# Home alone: A population neuroscience investigation of brain morphology substrates

**DOI:** 10.1101/2021.09.06.459185

**Authors:** MaryAnn Noonan, Chris Zajner, Danilo Bzdok

## Abstract

As a social species, ready exchange with peers is a pivotal asset - our “social capital”. Yet, single-person households have come to pervade metropolitan cities worldwide, with unknown consequences in the long run. Here, we systematically explore the morphological manifestations associated with singular living in ∼40,000 UK Biobank participants. The uncovered population-level signature spotlights the highly associative default mode network, in addition to findings such as in the amygdala central, cortical and corticoamygdaloid nuclei groups, as well as the hippocampal fimbria and dentate gyrus. Sex-stratified analyses revealed male-specific neural substrates, including somatomotor, saliency and visual systems, while female-specific neural substrates centred on the dorsomedial prefrontal cortex. In line with our demographic profiling results, the discovered neural imprint of living alone is potentially linked to alcohol and tobacco consumption, anxiety, sleep quality as well as daily TV watching. The secular trend for solitary living will require new answers from public-health decision makers.

## Introduction

Some animals have evolved by adapting to the benefits of living in a social group. In the primate lineage, this mode of living and coordination has probably improved the identification of scarce resources, and may have refined cooperating and dealing with predators and prey as a cohesive group (Dunbar & Shultz, 2017). As a result, various behaviour, neuronal, hormonal, cellular and genetic mechanisms have likely co-evolved to support these advantageous social forms (Robinson et al., 2008; Adolphs, 2009). For humans, the consequences of detachment from social group living can be expected to be pervasive due to the impoverished social environment. Indeed, social isolation is known to affect mental and physical well-being (Bzdok & Dunbar, 2020a). Such a state of deprived everyday stimulation is deemed so bad by society that it is used as an institutionalized form of punishment for individuals incarcerated in prisons (Cloud *et al*., 2015). Here we have investigated the impact of living alone on brain structure in a large community cohort of participants recruited from across the United Kingdom. This recently emerged population resource opens a unique window to see into the impact of the day-to-day social experience at the population scale in a naturalistic approach that goes beyond what traditional psychological and neuroscience experiments can do.

A wealth of neuroscience research now suggests that social abilities in humans and at least some non-human primates are realized by invoking a cohesive set of brain regions referred to as the ‘Social Brain’. Early support for the Social Brain idea came from evidence that, across species, the neocortex-to-brain volume ratio tracks the number of individuals per social group (Dunbar, 1992; Dunbar & Shultz, 2007a; b). This insight has been argued to imply that brain circuits particularly tuned to serving social processes have expanded via selection pressures acting over evolutionary time. For example, numerous subregions within the medial-temporal limbic system and medial prefrontal cortex show high neural responses to social information processing (e.g. face, expression, gaze) and dynamic social interaction (Noonan *et al*., 2016). This includes information of faces (Kanwisher *et al*., 1997; Ku *et al*., 2011), facial expression and gaze direction (Morin *et al*., 2015), species-specific vocalizations (Joly *et al*., 2012) and biological motion (Perrett *et al*., 1992). These brain circuits linked to social interplay are therefore key candidates in which differences in solitary living would be expected to manifest.

As such, we confront the question whether these recently evolved brain circuits that enable advanced coping with living in social groups may expose susceptibility when people undergo social scarcity in the environment. Clues to answer this question come from studies that have shown robust correlation of the size of individuals’ social network with indexes of structural and functional brain organisation. In humans, such studies have again typically implicated regions in the prefrontal cortex and the temporal lobe, particularly the amygdala (Bickart et al., 2011; Lewis et al., 2011; Von Der Heide et al., 2014; Noonan et al., 2018). Further, there is evidence that this pattern of effects may not simply reflect the individual’s predisposition towards seeking or avoiding social companionship. Instead, the brain may show plasticity effects in the face of recurring social experiences. In particular, Sallet and colleagues (Sallet et al., 2011) conducted controlled experiments with random allocation of monkeys to social housing for parallel laboratory studies (groups of 1-7 monkeys). This rare experimental feat demonstrated that the mid superior temporal sulcus (mSTS) and the medial prefrontal cortex both showed plasticity adaptations to daily living in a social group that has an experimentally imposed size. Later anatomical work has provided indicators that the temporal parietal junction (TPJ) is a strong candidate to be the human homologue of macaque mSTS (Mars *et al*., 2013), a region identified in humans as engaged in instantiating mental models of other people’s thoughts (Frith & Frith, 2006). These brain regions are also spatially contiguous with the default mode network (DNM) (Mars et al., 2012).

Collectively, these earlier studies bring to the surface how not only richness but also paucity of the social environment reverberates with specific brain systems. At its extreme, small-scale studies, in the context of arctic exploration or astronaut training and experience, have shown that enduring periods of social isolation is associated with increased stress hormone responses (Jacubowski *et al*., 2015; Weber *et al*., 2019). In the brain, these experiences of social isolation correlated with broad reductions in global cortical activity (Jacubowski *et al*., 2015; Weber *et al*., 2019) and specific reductions of the gray matter volume in prefrontal and hippocampal regions (Stahn *et al*., 2019). More generally, paucity of opportunity for social interaction in the real world has profound consequences for mental and physical health (Bzdok & Dunbar, 2020b). For example, social isolation is a major risk factor for age-related cognitive decline and Alzheimer’s dementia (Heinrich & Gullone, 2006).

Even the mere subjective perception of social disconnection from others, loneliness, takes a toll on mental health and cognition in all ages (reviewed by (Cacioppo & Hawkley, 2009). The perception of social disconnection is also associated with reduced overall life expectancy, and increases vulnerability to Alzheimer’s disease related dementias. Indeed, we recently identified brain signatures of loneliness in gray matter morphology, intrinsic functional coupling, and fiber tract microstructure and found that they converged on the DMN (Spreng et al., 2020). This study also identified brain signatures to be more pronounced in males than females. On its flipside, objective measures of social isolation have been linked to the limbic and salience networks (Schurz *et al*., 2021b). Again, there are sex-specific effects in the amygdala of various measures of social connection included not only household size, but also subject loneliness as well as objective social support (Kiesow *et al*., 2020). This array of robust brain-behavior associations speak to the relevance of social isolation has on the individual and the potential underlying neural substrates. As one possible interpretation, quantifiable sex-related divergences in social experience may be reflected in distinct neural profiles associated with living alone.

Finally, there are now swelling numbers of single-person households in numerous metropolitan cities across the globe. Hence, solitary living is becoming an increasing burden on modern societies (Raymo, 2015; Byron, 2019; Jackie Tang, 2019; Statistics, 2019). These compounding developments now warrant deeper understanding into the primary biology underlying lack of regular social interaction in the home environment. Decisive steps towards filling this knowledge gap may bring crucial insights into the associated mental and physical health consequences. In the present population neuroscience study, we take a naturalistic approach by utilising the large UK Biobank population imaging cohort (n=∼40,000 aged 40-69 years, mean age 54.9) to examine the gray matter correlates of living alone relative to living with other persons at home. We then explored putative sex-specific differences in the day-to-day experience of living alone, subsequently contextualized the results by their relation to perceived loneliness and regular social support, and conduct a careful demographic profiling analysis across key behavioral traits.

## Methods

### Population data source

The UK Biobank is a prospective epidemiology resource that offers extensive behavioral and demographic assessments, medical and cognitive measures, as well as biological samples in a cohort of ∼500,000 participants recruited from across Great Britain (https://www.ukbiobank.ac.uk/). This openly accessible population dataset aims to provide multimodal brain-imaging for ∼100,000 individuals, planned for completion in 2022. The present study was based on the recent data release from February 2020 that augmented brain scanning information to ∼40,000 participants. The present analyses were conducted under UK Biobank application number 25163. All participants provided informed consent. Further information on the consent procedure can be found elsewhere (http://biobank.ctsu.ox.ac.uk/crystal/field.cgi?id=200).

In an attempt to improve comparability and reproducibility, our study built on the uniform data preprocessing pipelines designed and carried out by FMRIB, Oxford University, UK (Alfaro-Almagro *et al*., 2018). Our study involved data from the ∼40,000 participant release with brain-imaging measures of gray matter morphology (T1-weighted MRI [sMRI]) from 48% men and 52% women, aged 40-69 years when recruited (mean age 55, standard deviation [SD] 7.5 years). Our study focused on single-person household status as a measure of richness of the social environment (Hawkley *et al*., 2003; Luhmann & Hawkley, 2016; Bzdok & Dunbar, 2020b). This self-reported item was based on the following question: “Including yourself, how many people are living together in your household? (Include those who usually live in the house such as students living away from home during term, partners in the armed forces or professions such as pilots)” (https://biobank.ndph.ox.ac.uk/showcase/field.cgi?id=709). Our analyses distinguished between people living by themselves (encoded as ‘1’) or living with other people (encoded as ‘0’) at home.

Binary target outcomes are found in widely used assessments of social embeddedness (Hawkley *et al*., 2005; Cyranowski *et al*., 2013). As one example, the Social Relationships scales of the NIH Toolbox (Cyranowski *et al*., 2013) feature the dimension of emotional social support. This dimension holds items such as “I have someone I trust to talk with about my problems”, or “I can get helpful advice from others when dealing with a problem”. A variety of studies showed such single-item measures of social traits to be reliable and valid (Mashek *et al*., 2007; Dollinger & Malmquist, 2009) Our own previous research has used yes-no items to study individuals who live alone.

### Multimodal brain-imaging and preprocessing procedures

Magnetic resonance imaging (MRI) scanners were matched at several dedicated imaging sites with the same acquisition protocols and standard Siemens 32-channel radiofrequency receiver head coils (3T Siemens Skyra). To protect the anonymity of the study participants, brain-imaging data were defaced and any sensitive meta-information was removed. Automated processing and quality control pipelines were deployed (Miller *et al*., 2016; Alfaro-Almagro *et al*., 2018). To improve homogeneity of the imaging data, noise was removed by means of 190 sensitivity features. This approach allowed for the reliable identification and exclusion of problematic brain scans, such as due to excessive head motion.

#### Structural MRI

The sMRI data were acquired as high-resolution T1-weighted images of brain anatomy using a 3D MPRAGE sequence at 1 mm isotropic resolution. Preprocessing included gradient distortion correction (GDC), field of view reduction using the Brain Extraction Tool and FLIRT (Jenkinson & Smith, 2001; Jenkinson *et al*., 2002), as well as non-linear registration to MNI152 standard space at 1 mm resolution using FNIRT (Andersson *et al*., 2007). To avoid unnecessary interpolation, all image transformations were estimated, combined and applied by a single interpolation step. Tissue-type segmentation into cerebrospinal fluid (CSF), gray matter (GM) and white matter (WM) was applied using FAST (FMRIB’s Automated Segmentation Tool, (Zhang *et al*., 2001)) to generate full bias-field-corrected images. SIENAX (Smith *et al*., 2002), in turn, was used to derive volumetric measures normalized for head sizes.

### Analysis of associations between living alone and gray matter variation

Neurobiologically interpretable measures of gray matter volume were extracted in all participants by summarizing whole-brain sMRI maps in Montreal Neurological Institute (MNI) reference space. This feature generation step was guided by the topographical brain region definitions of the widely used Schaefer-Yeo atlas comprising 100 parcels (Schaefer *et al*., 2018). The derived quantities of local gray matter morphology provided 100 average volume measures for each participant. The participant-level brain region volumes provided the input variables for our Bayesian hierarchical modeling approach (cf. below). As a data-cleaning step, inter-individual variation in brain region volumes that could be explained by variables of no interest were regressed out: age, age^2^, sex, sex*age, sex*age^2^, body mass index, head size, head motion during task-related brain scans, head motion during task-unrelated brain scans, head position and receiver coil in the scanner (x, y, and z), position of scanner table, as well as the acquisition site of the MRI data.

To examine population variation of our atlas regions in the context of household status, we purpose-designed a Bayesian hierarchical model, a natural choice of method building on our previous research (Bzdok *et al*., 2017a; Bzdok & Dunbar, 2020b; Kiesow *et al*., 2020; Kiesow *et al*., 2021; Schurz *et al*., 2021b). In contrast, classical linear regression combined with statistical significance testing would simply have provided p-values against the null hypothesis of no difference between participants living in a single-person household or not in each brain region. Instead of limiting our results and conclusions to strict categorical statements, each region being either relevant for differences in household size, our analytical strategy aimed at full probability distributions that expose how brain region volumes converge or diverge in their relation to household size as evidenced in the UK Biobank population. In a mathematically rigorous way, our approach estimated coherent, continuous estimates of uncertainty for each model parameter at play for its relevance in household situations. Our study thus addressed the question “How certain are we that a regional brain volume is divergent between individuals living alone or not?”. Our analysis did not ask “Is there a strict categorical difference in region volume between individuals living alone or not?”.

The elected Bayesian hierarchical framework also enabled simultaneous modelling of multiple organizational principles in one coherent estimation: i) *segregation* into separate brain regions and ii) *integration* of groups of brain regions in the form of spatially distributed brain networks. Two regions of the same atlas network are more likely to exhibit similar volume effects than two regions belonging to two separate brain networks. Each of the region definitions was pre-assigned to one of the seven large-scale network definitions in the Schaefer-Yeo atlas (Schaefer *et al*., 2018), providing a native multilevel structure to be modelled explicitly.

Setting up a hierarchical generative process enabled our analytical approach to borrow statistical strength between model parameters at the higher network level and those at the lower level of constituent brain regions. By virtue of exploiting such partial pooling of information, the brain region parameters were modelled themselves by the hyper-parameters of the hierarchical regression as a function of the network hierarchy to explain interindividual differences in solitary living. Assigning informative priors centered around zero provided an additional form of regularization by shrinking coefficients to zero in the absence of evidence to the contrary. We could thus provide fully probabilistic answers to questions about the morphological relevance of individual brain locations and distributed cortical networks by a joint varying-effects estimation that profited from several biologically meaningful sources of population variation.

Our model specification placed emphasis on careful inference of unique posterior distributions of parameters at the brain network level to discriminate individuals living with others (encoded as outcome 0) or those living alone (outcome 1) at their household:

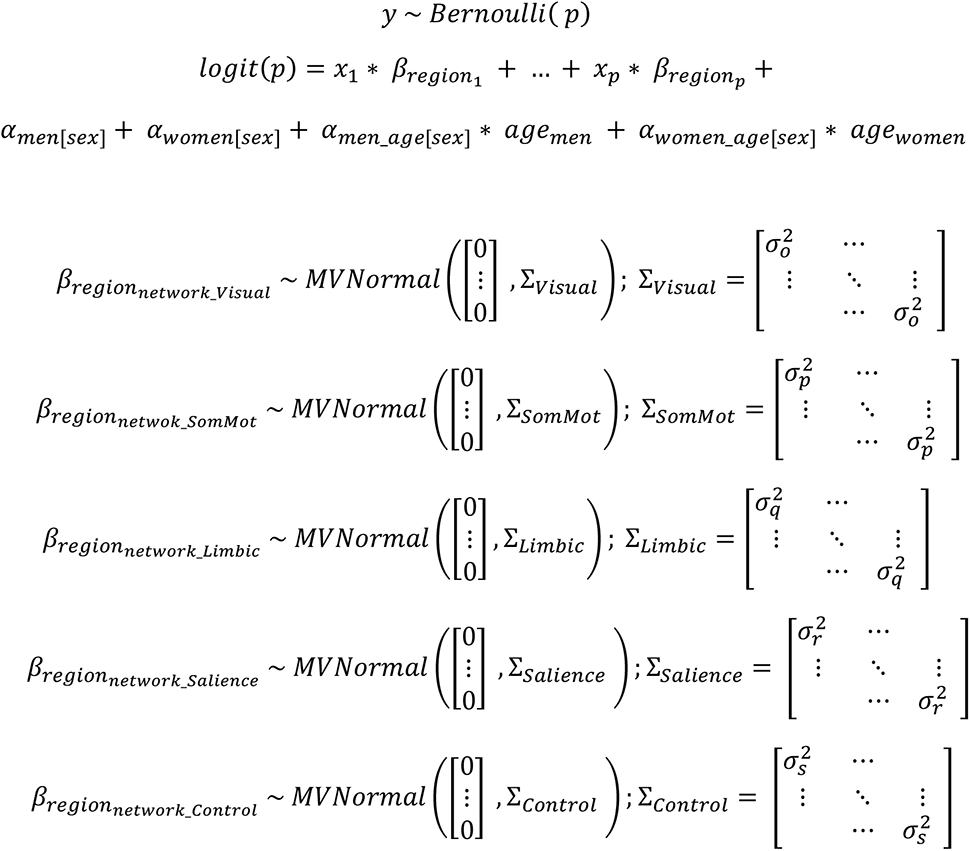

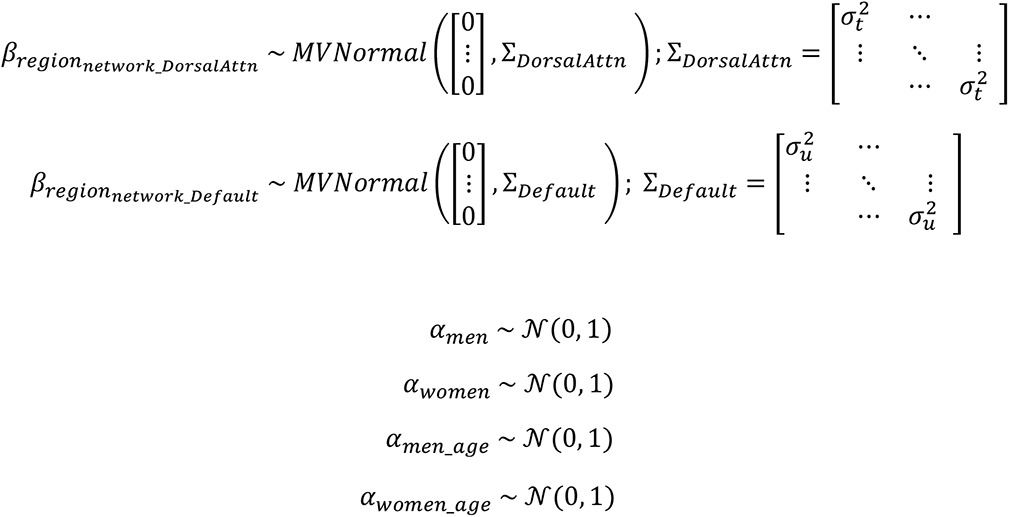

where *sigma* parameters estimated the overall variance across the *p* brain regions that belong to a given atlas network, independent of whether the volume effects of the respective constituent brain regions had positive or negative direction. As such, the network variance parameters *sigma* directly quantified the magnitude of intra-network coefficients, and thus the overall relevance of a given network in explaining lack of social interaction at home based on the dependent region morphology measures. All regions belonging to the same brain network shared the same variance parameter in the diagonal of the covariance matrix, while off-diagonal covariance relationships were zero.

Full probabilistic posterior distributions for all model parameters were inferred for the hierarchical modelling solution. By espousing a Bayesian attitude, we could thus simultaneously appreciate gray matter variation in segregated brain regions as well as in integrative brain networks in a population cohort. The approximation of the posterior distributions was carried out by the NUTS sampler (Gelman *et al*., 2014), a type of Markov chain Monte Carlo (MCMC), using the PyMC3 software (Salvatier *et al*., 2016). After tuning the sampler for 4,000 steps, we drew 1,000 samples from the joint posterior distribution over the full set of parameters in the model for analysis. Proper convergence was assessed by ensuring Rhat measures (Gelman *et al*., 2014) stayed below 1.02.

For illustration purposes, all brain images in MNI space were mapped onto a pial surface (Glasser *et al*., 2016) using the Connectome Workbench command-line tools.

### Post-hoc characterization of the brain substrates of solitary living regarding social isolation traits

Next, we sought to deepen insight into the set of relevant regions that was most robustly linked to residing in a single-person home. For this purpose, we quantified the strength of association of the volume measures from the top brain regions with external measures of objective and subjective social isolation that were not invoked in any previous steps of the analysis workflow: the opportunity of daily social exchange with others to confide is a well-accepted indicator for regular social support (data field: 2110; “How often are you able to confide in someone close to you?”) (Schurz *et al*., 2021a), while the experience of loneliness is commonly viewed to capture especially the feeling or personal impression of being social disconnected from others (data field: 2020; Do you often feel lonely? (Spreng *et al*., 2020)). Here, we examined all four possible combinations of these two complementary traits of social isolation in our UK Biobank sample. Given the four-group distinction setting, linear discriminant analysis was a natural choice of method. This classification machine learning algorithm (Bzdok, 2017; Bzdok *et al*., 2017b) afforded inferential statements about the effect sizes paired with the region-wise associations with each of the four disparate qualities of social isolation (i.e., each combination of subjective loneliness and objective social support).

### Demographic profiling analysis of the brain substrates of solitary living

We finally performed a profiling analysis of the brain regions that were most strongly associated with residing in a single-person home. We carried out a rigorous test for multivariate associations between our top region set and a diverse set of lifestyle indicators that exemplify the domains of a) basic demographics, b) personality features, c) substance-use behaviors, and d) social network properties (for details see https://www.ukbiobank.ac.uk/data-showcase/). Each of the behavioral variables and brain measures was z-scored across participants to conform to a mean of zero and a standard deviation of one. The brain variables that were submitted to this analysis were identified based on their importance in the context of solitary living (cf. above). These target brain regions were elected based on the (absolute) modes of the Bayesian posteriors of marginal parameter distributions at the region level.

Using the two separate variable sets, brain measurements and behavior measurements, we then carried out a bootstrap difference analysis of the collection of target traits in single-person versus multi-person households (Efron & Tibshirani, 1994). In 1,000 bootstrap iterations, we randomly pulled equally sized participant samples to perform a canonical correlation analysis (CCA), in parallel, according to household status (Miller *et al*., 2016; Wang *et al*., 2018). In each resampling iteration, this approach estimated the doubly multivariate correspondence between the brain and behavior indicators in each of the two groups. The ensuing canonical vectors of the leading CCA mode indicated the most explanatory demographic associations in a given pull of participants. To directly estimate the certainty of the brain-behavior cross-associations in the face of resample-to-resample variation, these canonical vectors of behavioral rankings, from CCA applications to single-person vs. multi-person households, were subtracted elementwise, recorded, and ultimately aggregated across the 1,000 bootstrap datasets.

We thus propagated the noise due to participant sampling variation into the computed uncertainty estimates of group differences in the UK Biobank population cohort. Statistically relevant behavioral dimensions were determined by whether the (two-sided) bootstrap confidence interval included zero or not in the 5/95% bootstrap interval. In a fully multivariate setting, our non-parametric modelling tactic directly quantified the statistical uncertainty of how a UK Biobank trait is differentially linked to brain-behavior correspondence as a function of household size.

## Results

### Network-level results whole population

By deploying an integrative Bayesian hierarchical modelling framework, we associated the objective experience of living alone with volume variation across the 100 brain regions that belong to the 7 spatially distributed brain networks that populate the human cerebral cortex, according to the Schaefer-Yeo reference atlas (Schaefer *et al*., 2018). At the network level, volume variation was most prominently associated with living alone in the default network, with the largest share of explained variance (posterior sigma=.065; 10-90% highest posterior density [HPD]=.044/.083; Fig. 1). The highest explanatory relevance of the collection of default network regions in living alone was followed by overall effects of the limbic network (sigma=.054, HPD=.001/.081), somatomotor network (sigma=.051, HPD=.019/.076), visual network (sigma=.050, HPD=.021/.074), as well as salience (sigma=.037, HPD=.002/.055), dorsal attention (sigma=.025, HPD=.001/.038), fronto-parietal (sigma=.021, HPD=.001/.031) networks. As such, our quantitative findings indicate the deepest layers of the neural processing hierarchy – the DMN regions – to play the strongest role in the brain manifestations of solitary living.

**Figure 1.**
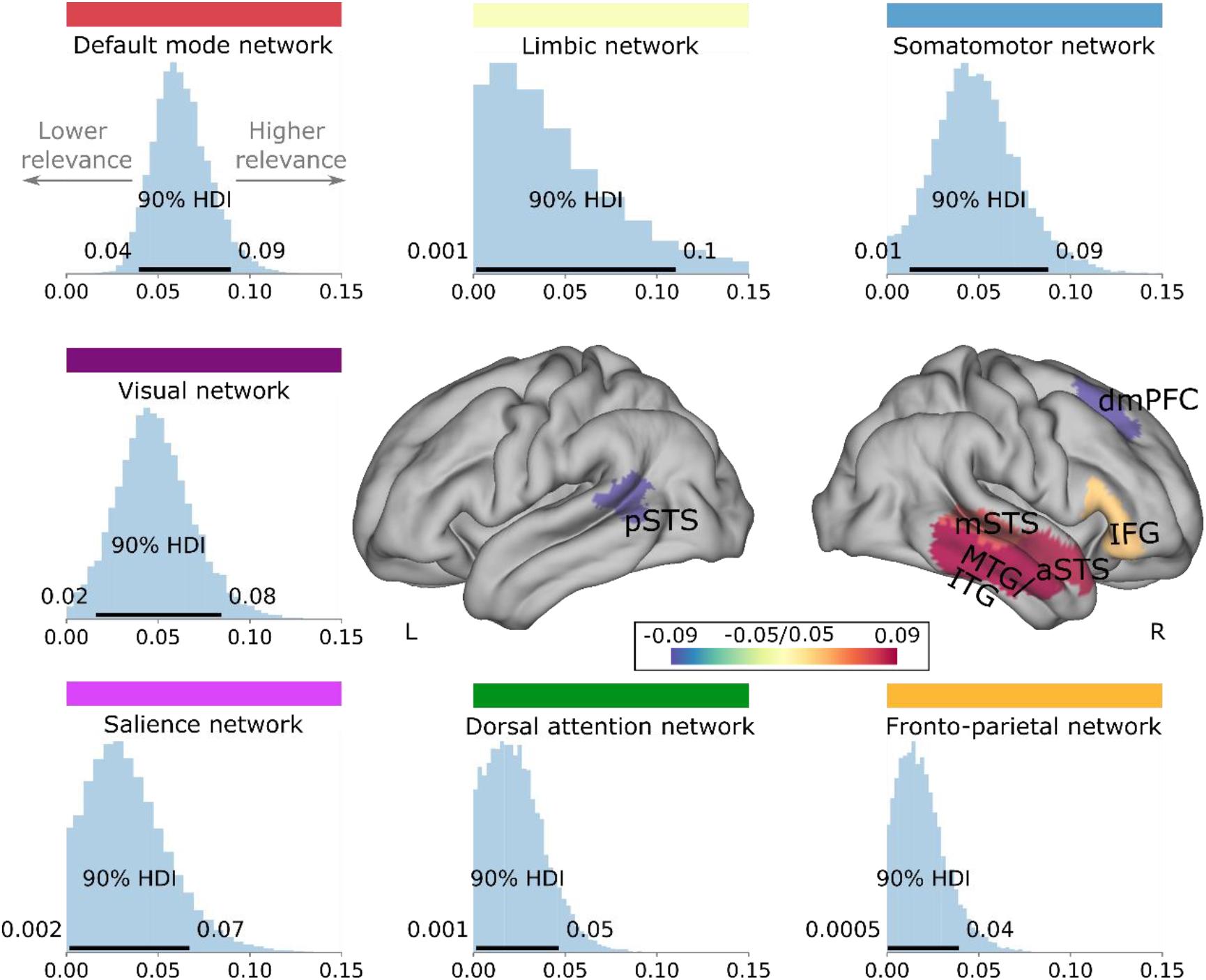
Solitary living is associated with default mode structure at the network and region level. Our Bayesian hierarchical modelling framework estimated the gray matter effects jointly of single regions and distributed networks of brain regions in explaining living alone. Roughly analogous to ANOVA, the network definitions could be viewed as factors and the region definitions could be viewed as continuous factor levels. Our framework allowed to begin quantifying the degree to which volume variation in each canonical network of regions reliably relates to living alone, as well as each separate region from those brain networks. Histograms show the inferred marginal posterior parameter distributions of the overall explanatory variance (sigma parameter) for each major brain network (volume measures in standard units). Horizontal black bars indicate the highest posterior density interval (HPI) of the model’s network variance parameters, ranging from 10 to 90% probability. Posterior distributions for the variance parameter (sigma) of each brain network are ordered from strongest (DMN; top left) to weakest (fronto-parietal; bottom right). The two brain renderings show the individual brain regions which were found to have the most robust relationship with living alone with their posterior parameter distributions (mean parameter). The brain regions that emerged as the most explanatory were in the lateral temporal lobe (pSTS, mSTS, aSTS, MTG/ITG), and frontal cortex (IFG, and dmPFC). a/m/pSTS = anterior/middle/posterior superior temporal sulcus, IFG = inferior frontal gyrus, dmPFC = dorsomedial prefrontal cortex, MTG = middle temporal gyrus, ITG = inferior temporal gyrus. L/R=left/right. Overall, the DMN and the limbic system showed overwhelming effects in explaining inter-individual variation in living alone.

Next, we investigated the impact of biological sex on the relationship between living alone and gray matter volume at the network-level. We found no salient differences in the degree of gray matter (GM) volume variation associated with living alone between the two sexes. Indeed, within the two groups the pattern of network effects were mostly similar to those of the whole population. For example, the top three networks that collectively explain most variance in women - limbic (posterior sigma=.072; 10-90% highest posterior density [HPD]=.002/.106; Fig. S1), somatomotor (sigma=.058; HPD=.015/.093) and default (sigma=.057; HPD=.032/.08) - were those that also collectively explained the most GM variation at the whole population, albeit in a different ranked order. Similarly, men showed significant GM variation within the DMN (sigma=.066; HPD=.037/.092) and limbic network (sigma=.063; HPD=.002/.096), but contrary to the whole population, the salience network (Ventral Attention; sigma=.061; HPD=.003/.087) explained the third most variance.

### Region-level whole population

We next inspected the inferred associations between living alone and regional brain structure. Using the previously described Bayesian hierarchical approach we focused on variation in GM volume in the 100 individual atlas regions (Fig. 1, Supplementary Table 1). Positive volume effects associated with living alone emerged in the right middle temporal gyrus/ inferior temporal gyrus (posterior mean=.089, 10-90% HPD=.033/.147), right anterior superior temporal sulcus (mean=.086, HPD=.031/.145), right middle superior temporal sulcus (mean=.080, HPD=.019/.136) and right inferior frontal gyrus (mean=.062, HPD=.015/.113). By contrast, negative volume effects became apparent in the left posterior superior temporal sulcus (mean=-.076, HPD=-.125/-.021) and right dorsomedial prefrontal cortex (mean=-.097, HPD=-.144/-.047). Lateral temporal subregions thus tended to explain the greatest amount of inter-individual variance in living alone.

### Region-level sex differences

When we turned to examine regional sex differences in the relationship between living alone and GM volume, we reported relevant effects in a range of association cortical regions broadly linked to action and perception. Our Bayesian hierarchical inference revealed a relatively right lateralized set of positive GM volume effects (Supplementary Table 2), indexing greater GM volume effects in men than women. These regions included the insula (mean=.076, 25-75% HPD=.013/.103), cuneus (mean=.075, HPD=.016/.099), precuneus/posterior cingulate cortex (mean=.059, HPD=.008/.099), motor/dorsal supplementary motor cortex (mean=.055, HPD=.006/.083) and posterior cingulate sulcus (mean=.052, HPD=.004/.079). Only motor/dorsal supplementary motor cortex showed positive volume effects in the left hemisphere (mean=.05, HPD=.003/.075). By contrast negative volume effects, indexing greater GM volume associations with living alone in women than men, were lateralized to the left hemisphere. The only significant effects evident were in the PFC; frontal polar cortex (mean=-.093, HPD=-.127/-.05) and dorsal premotor cortex (mean=-.07, HPD=-.109/-.02). These two frontal regions, both belonging to the DMN.

**Figure 2.**
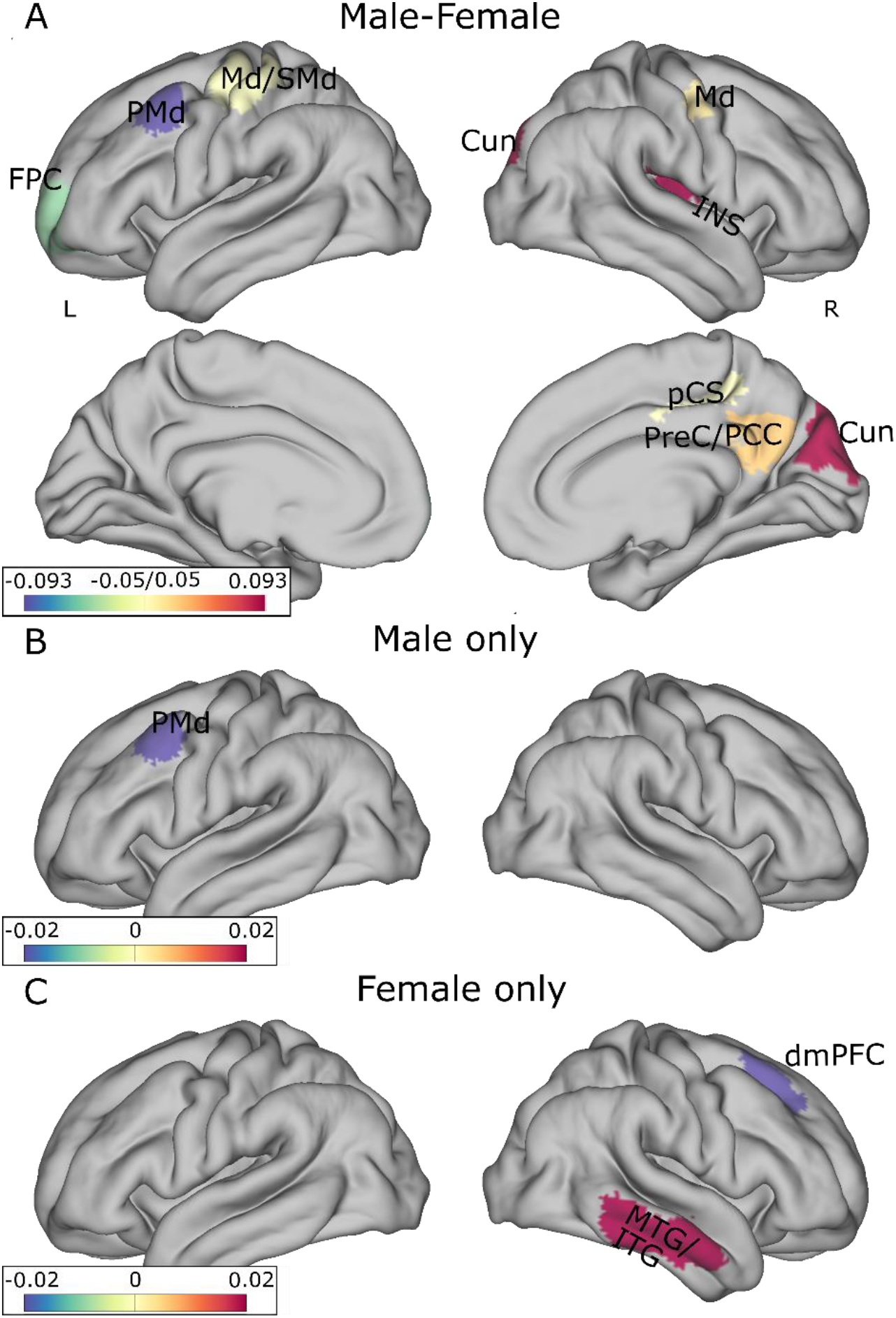
Degrees of sex bias characterize the gray matter substrates associated with living alone. Results highlight the brain regions that show different relationships to the experience of living alone in men and women. **A.** Sex contrast effects (male minus female) in the left (left column) and right (right column) hemispheres on lateral (upper rendering) and medial (lower rendering) at the region level (subtracting women’s posterior parameter distribution for a given effect from that inferred from males). For example, means of the posterior parameter distribution above zero can indicate a relatively male-biased effect with a positive volume effect associated with living alone (towards red color). Accordingly, in this case, for means below zero there would be a relatively female-biased volume effect for such brain-behavior association (towards blue color). **B-C.** Repetition of the Bayesian hierarchical analysis separately in (B) only males and (C) only females from our UK Biobank cohort: relevant gray matter effects (means of the marginal posterior parameter distributions). The neurostructural concomitants of living alone in men and women are notably different in a disparate assortment of brain regions. In men but not women, the dorsal premotor region emerges as robustly explanatory of living alone. Conversely the middle temporal gyrus/inferior temporal gyrus and dmPFC emerges in women but not men. FPC = frontal polar cortex, PMd = premotor dorsal, Md/SMd = dorsal motor/dorsal supplementary motor cortex, Cun = Cuneus, INS = insular cortex, pCS = posterior cingulate sulcus, PreC/PCC = PreCuneus/posterior cingulate cortex, MTG = middle temporal gyrus, ITG = inferior temporal gyrus, dmPFC = dorsomedial prefrontal cortex. L/R = left/right.

### Key regional effects are distinctly related to loneliness and quality of social support

The effects reported above describe brain regions that bear some type of association with living alone. But these effects by themselves cannot speak to whether these differences are related to felt or actual social isolation. In a sub-analysis, we thus addressed this question by teasing apart the contributions to GM volume effects with relation to subjective social isolation (loneliness) and/or objective social isolation (living alone). To this end we, conducted a post-hoc analysis in our UK Biobank sample that explicitly interrogated links to four participant subgroups: i) individuals who were not lonely and had good social support, ii) individuals who were lonely despite good social support, iii) individuals who were not lonely but had poor social support, and iv) individuals who were lonely and had poor social support. We focused the analysis on those six brain regions whose GM variation was found to explain relevant variation in living alone (Figure 3).

**Figure 3.**
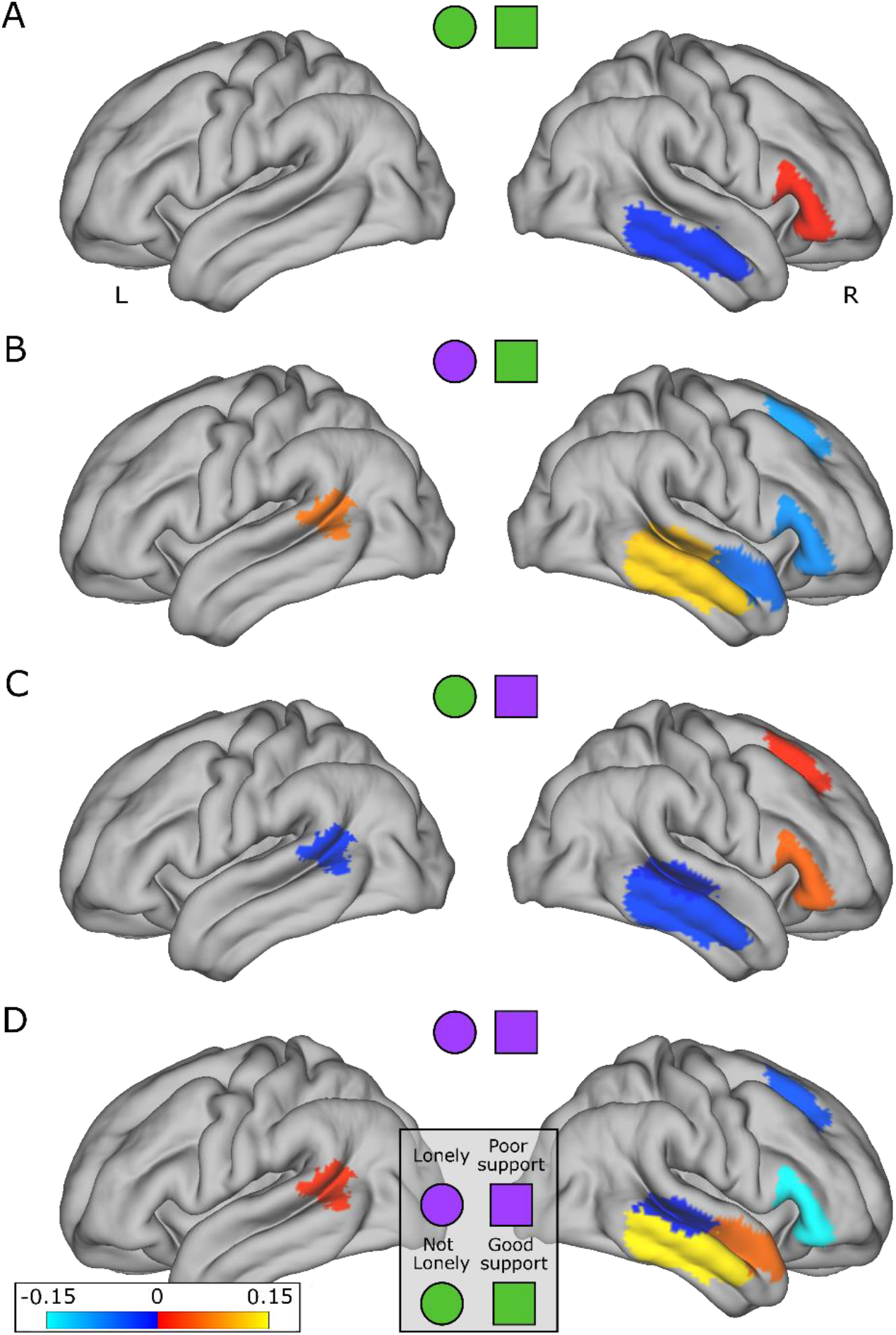
Top brain substrates of solitary living show differential links with objective and subjective isolation. As a means to further functionally annotate the brain correlates that we identified as relevant for solitary living (cf. Figure 1; Bayesian hierarchical model), we conducted a post-hoc analysis to examine the interindividual differences in volume variation in the identified relevant brain regions. This descriptive approach estimated how the participants of our UK Biobank sample can be distinguished based on their self-report measures of subjective and/or objective social isolation, that is, probing against all combinations of loneliness and social support. We thus aimed to map out which solitary-living correlates are preferentially linked to facets of social isolation that were external to the upstream analysis steps. Gray matter volume variation associated with individuals who do not feel lonely and indicate good social support (well surrounded, **A**). We show effects in individuals who report loneliness despite good social support (lonely, **B**), individuals who are not lonely despite poor social support (less social support, **C**), individuals who are lonely and have poor social support (lonely less social support, **D**). Effects were thresholded at 0.01 before surface mapping to the Connectome brain. Overall, the brain correlates of solitary living show especially strong volumetric relationships with loneliness. In particular, the MTG/ITG, IFG, and dmPFC are highlighted in loneliness.

Indeed, lonely individuals (regardless of quality of social support) consistently show GM effects across the network identified in the primary living alone analysis (B, D). This includes positive weights in the MTG/ITG regardless of the reported quality of social support. Notably, the anterior and posterior STS - both showing positive effects in individuals living alone - reverse sign in lonely individuals depending on quality of social support. For example, there was a positive pSTS and negative aSTS when social support structures are strong, and the opposite with poor social support (B vs D). The prefrontal regions both showed negative relationships with loneliness, regardless of social support (B, D). This was in line with the negative association between dmPFC GM and living alone, but opposite to the positive association between the IFG and living alone (Figure 1). By contrast, individuals who are well surrounded (not lonely), regardless of the quality of the social support network, generally show a complementary pattern of brain-behavior associations (A, C). Here the pattern of GM volume variation modulated by these two factors is predictably in the opposing directional weight than the living alone effects (Figure 1). For example, the temporal lobe clusters (particularly MTG/ITG) show positive effects in the whole population living alone analysis and negative effects when individuals are well surrounded (A, C). Similarly, the dmPFC cluster seen in individuals who are not lonely despite poor social support (C) showed reversed directional weight between original living alone contrast and the current analysis. However, this pattern is not always the case, with the IFG showing positive effects in individuals who live alone and individuals who report not being lonely (regardless of social support quality A,C).

### Amygdala nuclei gray matter relationships with living alone

We next turned to a fine-scale assessment of a closely associated subcortical structure with our identified whole-brain correlates of solitary living - the amygdala. To this end, we examined the relationship between amygdala nuclei GM and living alone using Bayesian inference (Figure 4, Supplementary Table 3). Notably, there was a strong lateralisation in the amygdala volume effects. Further, all but one amygdala subregion with a relevant effect showed a negative association between GM volume and living alone. Negative volume effects associated with living alone emerged in the right central (posterior mean=-.048, 10-90% HPD=-.078/-.017), right cortical (mean=-.057, HPD=-.095/- .019), right cortico-amygdaloid-transition (mean=.08, HPD=.019/.136), and left accessory basal nucleus (mean=-.109, HPD=-.196/-.018). The left accessory basal nucleus was the only robust effect we observed in the left amygdala. Conversely, we observed a positive effect in the right accessory basal nucleus (mean=.224, HPD=.136/.319). In sum, the majority of the salient relationships between amygdala nuclei GM volume and solitary living were on the right hemisphere. Most of these effects were also negative amygdala-household-living associations.

**Figure 4.**
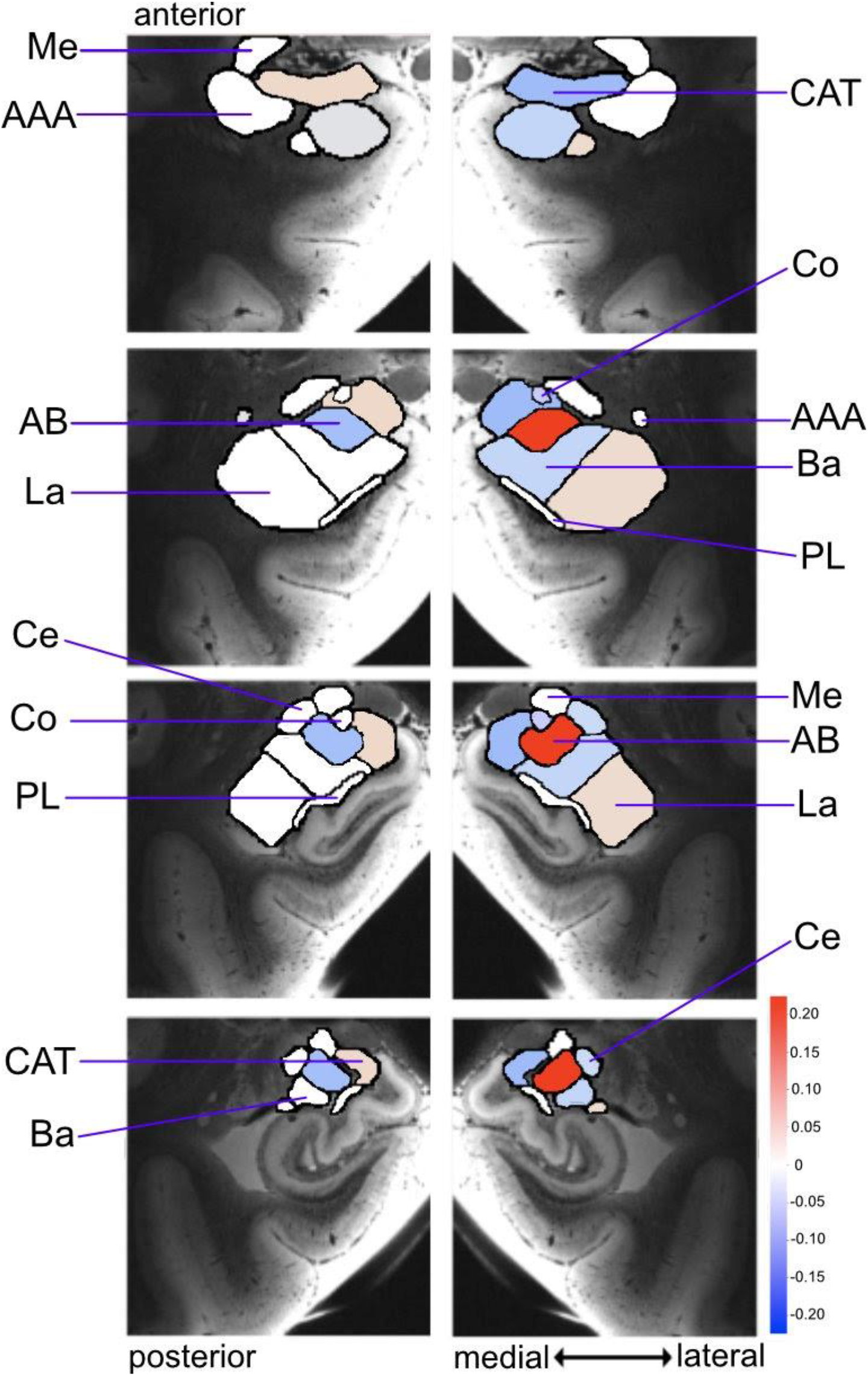
Specific amygdala nuclei groups are differentially affected in solitary living. Shows the results from Bayesian modelling applied to the amygdala based on the 9 cytoarchitectonically distinguishable nuclei groups from an automatically derived amygdala segmentation protocol (Saygin, Klienmann et al. 2017). The inferred Bayesian posterior parameter distributions indicate where volume variation can explain single person households. Shown as means of the marginal posterior parameter distributions, the results are mapped to 4 consecutive coronal sections of the left and right amygdala from anterior (top) to posterior (bottom) (hot/cold colors = positive/negative volume associations). The left and right accessory basal nucleus show particularly strong, but opposing volumetric relationships. ME = Medial, AAA = Anterior Amygdala Area, CAT = Cortico-amygdaloid Transition Area, Co = Cortical, AB = Accessory Basal, La = Lateral Nucleus, Ba = Basal Nucleus, Ce = Central, PL = Paralaminar Nucleus. Overall, the amygdala nuclei with strong relationships to solitary living were primarily in the right hemisphere.

### Sex differentiation in amygdala nuclei relationships with solitary living

Given identified sex deviation in the association of cortical regions with living alone, we then sought to examine possible sex differences in amygdala nuclei (male – female, Supplementary Table 4). For example, male-biased positive volume effects can be indexed by greater volume contributions in men than women with less regular social interaction at home. Positive volume effects (that can be indicative of larger GM volume effect in men than women) were identified in the right paralaminar (mean=.084, 25-75% highest posterior density [HPD]=.040/.139), and right central nuclei (mean=.039, HPD=.002/.068), as well as the left lateral nucleus (mean=.091, HPD=.037/.125), and left anterior amygdaloid area (mean=.051, HPD=.021/.085). Negative sex-biased volume effects (that can be indicative of larger GM volume effect in women than men) were evident in right cortical (mean=-.09, HPD=-.124/-.045) and right lateral nuclei (mean=-.140, HPD=-.181/-.087), in addition to the left central nucleus (mean=-.067, HPD=-.096/-.034). We thus found various amygdala nuclei which showed diverging sex effects with respect to living alone. The sex effects also showed opposite patterns for the left and right amygdala for some nuclei. For example, the right central nucleus and left lateral nucleus showed a relationship of greater volume in men than women, while their opposite hemisphere counterparts, the left central nucleus and right lateral nucleus, showed greater volume effects in women than men.

### Hippocampus subregion volumetric relationships with living alone

Next, we examined the variation in volume amongst hippocampal subregions that explain the trait of living alone (Figure. 5, Supplementary Table 5). Several anatomical subregions in the hippocampus head showed relevant volume effects for the target phenotype. For example, our fine-resolution mapping approach identified positive volume effects in the head of the hippocampus in left CA1 (mean=.048, 10-90% highest posterior density [HPD]=.003/.094), right molecular layer (mean=.040, HPD=.008/.071), bilateral presubiculum (right mean=.043, 10-90% HPD=.002/.084, and left mean=.054, HPD=.014/.096), right CA2/3 (mean=-.09, HPD=-.127/-.054), and right dentate gyrus (mean=.099 HPD=0/0.203). Additionally, we identified salient negative volume effects in the head in the left dentate gyrus (mean=-.034, HPD=-.064/-.003) and left molecular layer (mean=-.034, HPD=- .064/-.003). We also found relevant volume effects in the body of the hippocampus, including the bilateral presubiculum (right mean=.046, HPD=.013/.08, and left mean=-.044, HPD=-.079/-.01), right dentate gyrus (mean=-.113, HPD=-.192/-.038), and left CA4 (mean=.094, HPD=.018/.175). Overall, our model pinpointed various robust relationships between the hippocampus at a subregion resolution and living alone, many of which were located towards the anterior (head) portion of the hippocampus.

**Figure 5.**
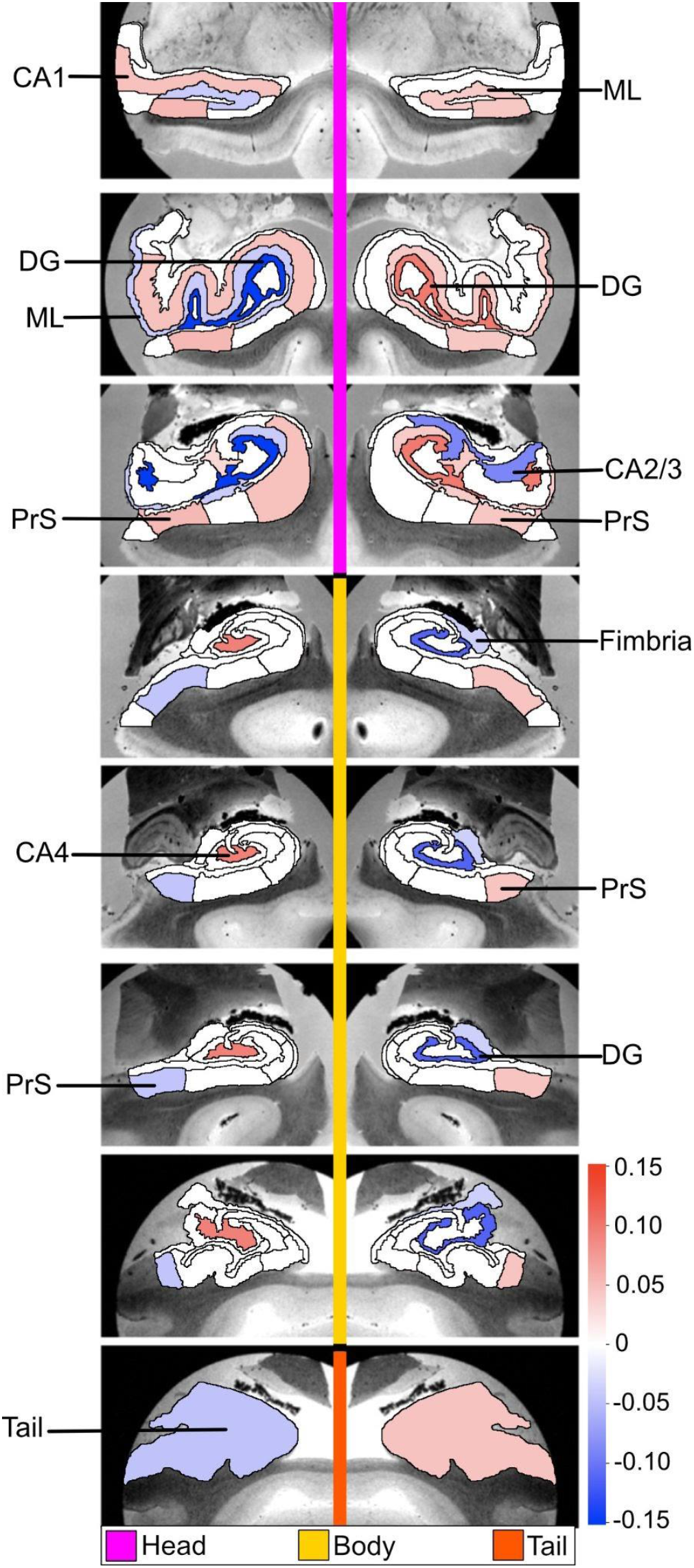
Specific hippocampus subfields are differentially affected by solitary living. The hippocampus subregions have robust links to living alone, indicated by our Bayesian model inference based on 38 subregions from an automatically derived hippocampus MRI image segmentation protocol (Iglesias, Augustinack et al. 2015). Shown as means of the marginal posterior parameter distributions, volume variation that can be explained by single person households by each specific hippocampus subregion mapped onto 8 consecutive coronal sections of the left and right hippocampus from anterior (top) to posterior (bottom) direction (hot/cold colors = positive/negative volume associations). The majority of subregions with robust effects were additionally located towards the head portion of the hippocampus. CA = cornu ammonis, PrS = presubiculum, GC = granule cell layer of dentate gyrus, DG = dentate gyrus, ML = molecular layer. Together, the subregions which explain inter-individual variation in living alone tend to have opposite effects in the left and right hippocampi.

In addition to the general trend of stronger structural associations of living with features of the head of the hippocampus than the body, we observed varying patterns of bilateral and lateralized volume effects. For example, the right and left presubiculum head both showed strong positive effects. However, the other laterality patterns show the opposite direction of effects comparing the two hemispheres. For example, the presubiculum body showed positive volume effects on the right and negative effects on the left. A similar pattern is found in the hippocampal tail (right mean=.046, 10-90% highest posterior density [HPD]=.016/.079, left mean=-.046, HPD=-.079/-.015), dentate gyrus head, and molecular layer head. Unilateral positive volume effects were additionally found in the left CA4 body and left CA1 head. In the right hemisphere, unilateral negative volume effects were found in the fimbria (mean=-.036, HPD=-.059/-.014), CA2/3 head, as well as the dentate gyrus body. As such, there were generally diverging relationships between right and left hippocampal subregion volumes and solitary living.

### Sex differentiation in hippocampus subregion relationships with solitary living

Sex-specific analyses of hippocampal subregions (male – female, Supplementary Table 6) revealed positive volume effects in the right GC-ML-DG-head (mean=.137, 25-75% highest posterior density [HPD]=.029/.241), right CA4 body (mean=.133, HPD=.036/.215), right parasubiculum (mean=.01, HPD=.076/.128), and the right hippocampal fissure (mean=.063, HPD=.032/.092). We also identified two subregions in the left hemisphere with greater GM volume effects in men than women: molecular layer body (mean=.113, HPD=.08/.14) and the fimbria (mean=.043, HPD=.017/.069). By contrast, negative volume effects (which can indicate greater GM effects in women compared to men) were found in the right molecular layer body (mean=-.047, HPD=-.076/-.017), right hippocampal tail (mean=-.066, HPD=-.098/-.034), right subiculum body (mean=-.086, HPD=-.127/-.048), and right CA2/3 head (mean=-.117, HPD=-.157/-.08). There were also a number of subregions in the left hemisphere with greater GM volume effects in women than men: parasubiculum (mean=-.027, HPD=- .055/-.003), presubiculum body (mean=-.054, HPD=-.095/-.022), and CA4 head (mean=-.12, HPD=- .23/-.018). Overall, we isolated a collection of hippocampal subregions that featured robust incongruencies of structural relationships with living alone depending on sex. Right and left hippocampal subregions which also showed differential associations based on sex included the parasubiculum and molecular layer body. For example, the right parasubiculum and left molecular layer body showed greater volume in men than women, while the left parasubiculum and right molecular layer body showed greater volume in women than men.

### Self-medicative and protective factors linked to living alone are pinpointed by demographic profiling

Finally, we performed a demographic profiling analysis that set out from the brain regions that were most strongly associated with living alone (Figure 1). We tested for multivariate cross-associations between these regions (see methods) and a diverse set of factors that covered the domains of (a) basic demographics, (b) personality features, (c) substance-use behaviors, and (d) social network properties. The resulting associations revealed that the largest real-world explanatory factors (but most variance) that accounted for the GM volume effects were linked to interindividual differences in everyday behaviour. These lifestyle indicators included self-medicative behaviours such as time spent watching television, past smoking frequency, alcohol intake on a typical day drinking day, alcohol intake frequency. Notably, potentially protective social factors also showed a strong association with the identified top brain regions, specifically the number of sisters and the number of brothers, suggesting family structure plays an important role in the social support system of individuals in a single-person household.

**Figure 6.**
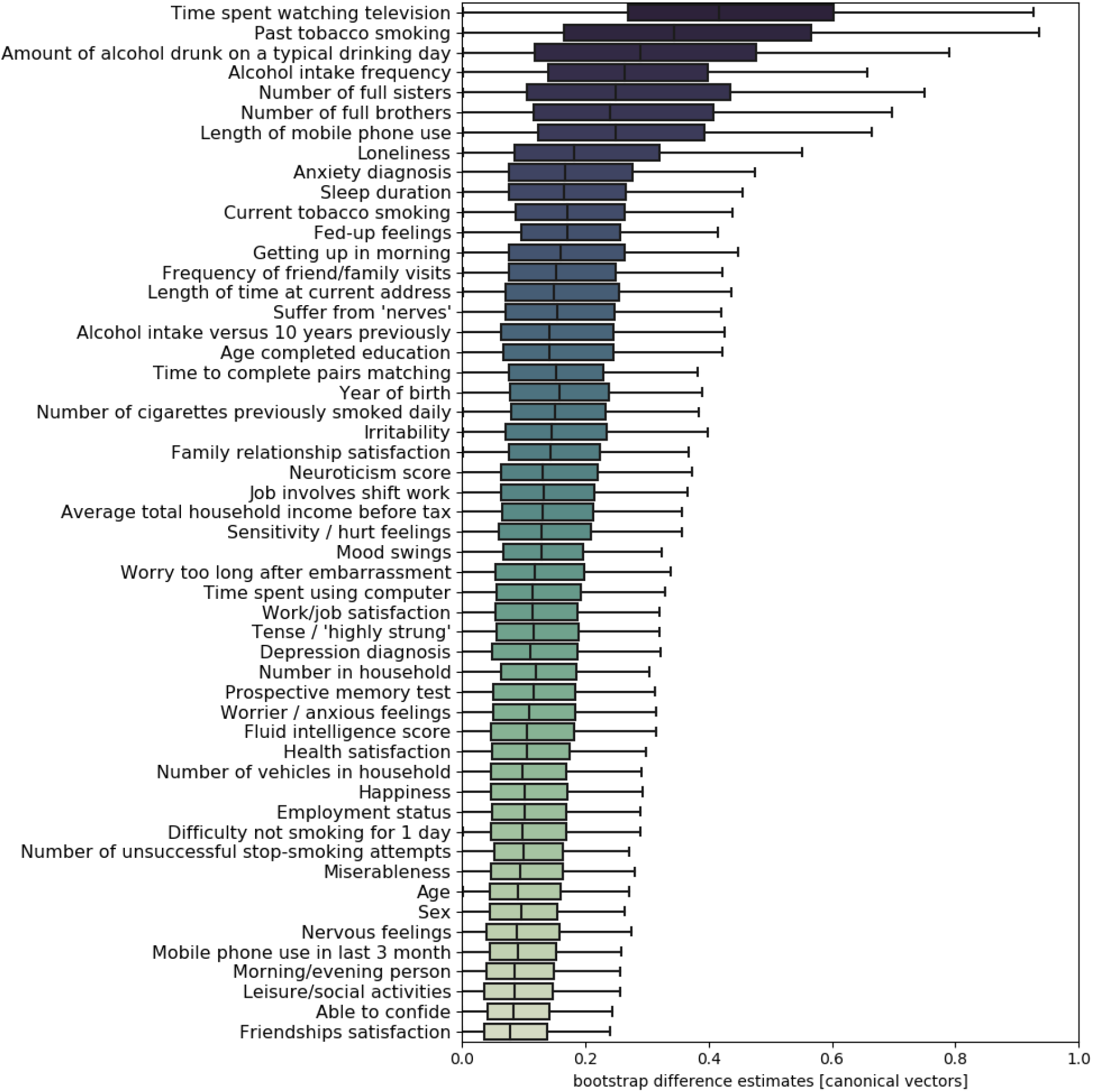
Demographic profiling analysis ranks lifestyle factors by relation to solitary living substrates in the brain. Multivariate pattern-learning (cf. Methods) was used to explore how the top brain regions (see Fig. 1) are linked to a portfolio of behavioral indicators in individuals living alone or with others. Behavioral markers covered domains of mental and physical well-being, lifestyle choices, and social embeddedness. In 1,000 bootstrap resampling iterations, our entire pattern-learning pipeline was repeated separately in the two participant groups: UK Biobank participants who live alone vs. with others. The computed differences in modelled brain-behavior associations between both groups (i.e., diverging canonical vector entries) were gathered across the 1,000 perturbed re-draws of our original sample to obtain faithful bootstrap intervals. The derived estimates of uncertainty directly quantified how group-related deviations vary in the wider population. The boxplot whiskers show the interquartile range (i.e., 25/75% interquartile distance derived from bootstrap resampling distributions). The highlighted divergences in individuals living in a single-person household reveal characteristics of these population strata that implicate indicators of media consumption, health and smoking behavior, as well as alcohol consumption at the population level.

## Discussion

Single-person households are becoming more common around the world, especially in many metropolitan cities (Nations, 2019). This unprecedented circumstance reduces the amount of daily social exchange for many people, with measurable sequelae for brain and behavior. Despite the known mental and physical health costs of solitary living, there is a knowledge gap in our understanding of the relationship between living alone and the brain at the population level. To begin addressing this need, the present population neuroscience study set out to systematically trace out brain manifestations linked to living alone in the ∼40,000 UK Biobank cohort. We uncover a population-level signature that highlights structural alterations in the highly associative DMN, in addition to subregion-specific effects in particular hippocampus subfields and amygdala nuclei. Sex-specific effects emerged in the highest association circuits in medial prefrontal cortex of women. Instead, males showed these effects at the intermediate (salience network) and lower (visual and somatomotor cortex) layers of the neural processing hierarchy.

Our study showed the DMN to yield broad network-wide associations with living alone. Additionally, at the region-level, our analysis uncovered that all regional GM effects that distinguish single-household individuals from those living with one or more persons at home coalesce to parts of the DMN. Specifically, we identified robust GM volume effects in individuals living alone in the pSTS and dmPFC. Additionally, we identified consistent gray matter volume effects in a number of superior and middle temporal lobe regions in individuals living alone. Based on several decades of social neuroscience research, the DMN is well known to typically show neural activity responses during tasks in the social domain including perspective taking capacities (Theory of Mind) as well as certain forms of empathy (Frith & Frith, 2006). Furthermore, DMN aberration is at the cross-roads of a number of neurological and psychiatric conditions with aspects of disordered social cognition, such as Alzheimer’s disease (Hafkemeijer *et al*., 2012) and autism (Anderson *et al*., 2011). It is therefore intriguing that, while our analysis approach was not specifically tuned to a particular brain system, the DMN emerged as a central point of convergence in solitary living across analyses.

Our collective findings not only highlight the DMN in understanding the sociocognitive factors associated with living alone, but replicate key findings from experimental studies in macaque monkeys. Sallet and colleagues (2011) showed associations of the GM volume with the size of an individual monkeys’ social group size, defined as the number of animals the individual shared their home cage with, ranging from single-housed to seven-socially housed animals. In the present study in humans, we report a negative relationship between GM and living alone in the dmPFC and the posterior STS. Our human dmPFC atlas region overlaps with the likely human homologue of the monkey area 9/46D, identified by Sallet et al. as larger in monkeys living in larger social groups. In humans, this region is associated with neural activity responses when predictions are made and updated about the intentions of others (Behrens *et al*., 2008; Seo & Lee, 2008). By contrast, the pSTS atlas region, like the dmPFC, is believed to be involved in theory of mind. This part of the temporal lobe is dorsally adjacent to a possible candidate of the human homologue of the mSTS identified in the macaque as larger in animals living in larger social groups; the posterior temporal parietal junction (Mars *et al*., 2013). Given evidence that dmPFC neurons are involved in predicting another social agent’s choice (Seo *et al*., 2014), one possible interpretation of the collective present and previous findings in macaques and humans is that these kinds of social predictions occur less frequently in the absence of social interactions and result in reduced gray matter in individuals who live alone.

Living alone is associated with detrimental physical and emotional consequences (Bzdok & Dunbar, 2020b). Many people know from their own experience that one does not need to physically be alone to succumb to the subjective feeling of loneliness. The results from the present study reveal the neurobiological commonalities between the emotional states of loneliness and living alone in the UK Biobank cohort. We find that many of the regions associated with single-person households also bear some relation with loneliness. First, our findings speak to previous work showing positive association between MTG/ITG and pSTS and the experience of loneliness (Spreng *et al*., 2020) but now observe this relationship is linked to the quality of social support. For example, there is a main effect of loneliness in MTG/ITG GM effects, regardless of the quality of social support, with the predicted positive GM effects between MTG/ITG and loneliness in individuals who report good and poor social support systems. By contrast, we found associations between loneliness and social support quality in pSTS GM effects. GM effects are positive when loneliness is reported in individuals who have good social support, but negative when individuals have a poor social support system, regardless of their sense of loneliness. Further, we did not always observe a corresponding association between living alone and the state of loneliness, that was seen in the MTG/ITG (i.e., positive GM effects in living alone, positive GM effects in lonely states, negative GM effects in well surrounded individuals). For example, the IFG showed the counterintuitive pattern; positive effects in individuals living alone, but negative GM effects in lonely individuals and positive GM effects in well surrounded individuals. Future research should examine the interaction between these three social factors and link neural relationships with behaviour to further identify adaptive and maladaptive effects.

An important source of interindividual variability in living alone turned out to be sex in our present study. In the UK, more men live alone before the age of 65 years, but notably this pattern reverses after that age (Esteve *et al*., 2020). Given known sex bias in primate behavioural and social development (Baron-Cohen *et al*., 1999; Key & Ross, 1999; Silk *et al*., 2003; Bhattacharya *et al*., 2016; Amici *et al*., 2019; Amici & Widdig, 2019), and various sex-dependent neuroanatomical differences reported in the amygdala, hippocampus, and various cortical regions (Lenroot & Giedd, 2010; Ritchie *et al*., 2018; Kiesow *et al*., 2020), we examined sex differences in neurobiological variability to solitary living. The sex-focused analyses corroborated the findings from the full sample, but notable patterns of differences became apparent between the sexes. Living alone in men was associated with a stronger negative volume effect than women in the frontal cortex and especially its medial portion, a region associated with tracking the significance of multiple goals in parallel, as well as switching between them (Boorman *et al*., 2009). By contrast, living alone in women was associated with more negative GM relationships, compared to men, in a number of visual, sensory motor and attentional regions, as well as relatively posterior subregions within the DMN including precuneus and cingulate gyrus. In fact, the reported effects did not localise to the higher-order association areas but to regions known to be involved in perception, memory and action, which may reflect evidence of sex differences in cognitive abilities (Asperholm *et al*., 2019). For example, sex differences in face processing, such as women judging faces as more positive and arousing than men, may translate to fundamental differences in lower-level perceptual experiences of men and women who live alone (Lewin & Herlitz, 2002; Proverbio, 2017; Mishra *et al*., 2019; Olderbak *et al*., 2019). Future research will be needed to directly link biological and cognitive differences to the sex-specific differences in behavioural strategies adopted when living alone. For example, women tend to entertain larger social networks and maintain more close friendships than men, especially later in life (Dunbar, 2018). This observation may act to protect women from the negative elements of living alone. By contrast, solitary living men, in this cohort, may be particularly adversely affected after retirement age if their social circles are grounded in their working environment.

In our pattern analyses dedicated to the amygdala at subregion resolution, living alone was associated with distinct anatomically defined nuclei groups. Besides bilateral effects in the accessory-basal nuclei group, the central nuclei group, cortical nuclei group and corticoamygdaloid transition all showed effects preferentially in the right hemisphere. The laterobasal nuclei group is commonly conceptualized as a likely integrator of preprocessed visual, auditory, gustatory, somatosensory, and, in part, olfactory environmental information (Aggleton *et al*., 1980; Iwai & Yukie, 1987; Stefanacci & Amaral, 2002; Yukie, 2002). As such, living alone may relate to stimulus-value associations subserved by the human laterobasal nuclei group that are believed to be implicated in associative processing of environmental information and the integration with self-relevant cognition in a way that is biased to the right brain hemisphere. Instead, the centromedial nulcei group, the amygdala’s putative major output center, has been related to integration of information originating from various intra-amygdala circuits to mediate behavioral and autonomic responses (Pessoa & Adolphs, 2010), including motor behavior and response preparation in humans. These amygdala subregion deviations may in part reflect the previous observation that socially deprived individuals show worse aptitude at significance detection, such as in detecting social cues from other’s faces or gestures to be overly negative and allocating attentional resources accordingly (Cacioppo *et al*., 2009). Such individuals are also known to react differently towards others, such as part of approach-vs-avoidance decisions and facial motor responses (Cacioppo *et al*., 2009). A right-hemispheric bias in such stimulus-response cycles could be related to the previous neuroimaging observation that the right hemisphere shows attention- and stress-related differences in socially deprived individuals (Cacioppo & Hawkley, 2009). Indeed, it has been proposed that social disconnection may trigger an evolutionary alarm signal and effects appear to lateralise to the right hemisphere as it could reflect increased attention towards threat and may link to the right ventral attention stream (Eisenberger *et al*., 2003; Cacioppo & Hawkley, 2009); flagging survival-relevant information in the environment.

Indeed, both the amygdala *and* hippocampus are known to be affected by social stress. This includes long-term changes in gross morphology, dendritic remodelling (retraction in CA1 and CA3 in the hippocampus and expansion in amygdala), functional connectivity and changes in neurogenesis (Woolley *et al*., 1990b; Watanabe *et al*., 1992b; Magarin & McEwen, 1995; Magariños *et al*., 1996; Magariños *et al*., 1997b; Vyas *et al*., 2002; Vyas *et al*., 2006; Anacker *et al*., 2018; Biggio *et al*., 2019a). Here we show robust associations between living alone and GM structure at hippocampal subfield scale). This resolution goes far beyond previous studies, which were often limited to a crude posterior/anterior division in the primate brain (Fanselow & Dong, 2010). We report numerous bilaterally coherent effects in the molecular layer head, pre-subiculum, para-subiculum and hippocampus tail, many of which have been modulated by social experience and stress. Social isolation is an extreme stress trigger and when induced by long term confinement is associated with broad reductions in global cortical activity and increased cortisol levels (Jacubowski *et al*., 2015; Weber *et al*., 2019). While inherently methodologically and ethically challenging to manipulate, findings from a number of studies in this area align with the current results. For example, experimentally induced social isolation during adolescence in monkeys chronically alters functional connectivity between the hippocampus and amygdala, and frontal cortical structures (Yuan *et al*., 2021). Similarly, in humans, the relative social isolation induced by a 14 month expedition of the Antarctic was exploited by researchers to reveal decreased GM volume in the DG hippocampal subfield, and decreased markers of neurogenesis at the end of the expedition (Stahn *et al*., 2019). Our observed effects in the DG (including the left CA4 body, right GC-DG-ML body, and bilateral GC-DG-ML head) fit with these and other studies of the neurostructural concomitants of living in limited social environments (Gould *et al*., 1998; Kempermann *et al*., 1998; Stranahan *et al*., 2006; Ibi *et al*., 2008; Dranovsky & Leonardo, 2012; Li *et al*., 2013; Anacker *et al*., 2018; Biggio *et al*., 2019b). The anterior portion of the DG/hippocampus has also been found to be particularly associated with stress susceptibility (Anacker *et al*., 2018). The chronically stressful experience of solitary living may thus manifest in the form of altered DG structure and function, but also in cooperating anterior (head) structures. For example, we found robust effects in the molecular layer head in both the right and left molecular layer - a region which has been well-described as being particularly sensitive to chronic stress (Gould *et al*., 1990; Woolley *et al*., 1990a; Watanabe *et al*., 1992a; Magariños *et al*., 1997a). Finally, we also bilaterally identified both presubiculum subfields (head, body) in the context of solitary living. The presubiculum is composed of grid cells (Boccara *et al*., 2010) and recent work has suggested the hippocampus tracks social relationships in the form of a social cognitive map that relies on a hexagonal coding structure (Tavares *et al*., 2015). A strong relationship between the presubiculum and living alone may therefore indicate an alteration of the neural underpinnings of a robust cognitive map of social spaces.

Finally, we charted brain-behavior associations between explanatory real-world factors and variation in the set of brain regions associated with living alone. At population level, this test for robust cross-associations suggest that one set of factors, such as smoking and frequent alcohol intake, may reflect compensatory or self-medicating associative behaviours that run parallel to living alone. By contrast, family structure, indexed by numbers of brothers and sisters may speak to a protective role linked to the discovered brain-behavior cross-associations. This insight may reflect the stable nature of a sibling relationship, compared to friendship circles which may be periodically disconnected. Further, small but significant variance in GM effects were also explained by individual differences in loneliness and the ability to confide a social support structure, which fits well with the analysis that explored the interactions between these three factors. Indeed we can see evidence that many of these behavioural factors have changed at the population level during this period of social restrictions during the pandemic, with increased total video viewing time (including TV and online streaming, (OfCom, 2020)), increased intake of alcohol in UK samples, particularly women (Sallie *et al*., 2020; Jackson *et al*., 2021), and increased smoking, mostly in younger age groups (Jackson *et al*., 2021). Collectively, these behavioural factors, including the likely positive effect of siblings, should be studied carefully alongside future investigations into the impact of living alone as they could provide targets to support individuals in such social environments.

For millennia, primates have socially cohabited. However, it is only over the last 10-20 years that we have seen a significant trend for more people to live alone and to reside at a greater geographically distant from their immediate families. The parallel increase in the frequency of global crises also acts to accelerate and aggravate the progressive dislocation and alienation of normal social forms of living. At the extreme, and as a result of the coronavirus pandemic, there was more than 50% of the world’s entire population under stay and home orders in April 2020. These unusual global circumstances and other extraordinary events, such as natural catastrophes or abrupt economic change, are likely to disproportionately jeopardize the well-being of people who live alone, increasing demands on both individual resilience but also financially on government and charity resources in the future. While online social networks can partially recapitulate real-world networks (Kanai *et al*., 2012; Dunbar, 2016) they cannot replace them. Consequently, a growing appreciation of cognitive, psychological and neural implications of solitary living and loneliness could directly inform social and health policies.

## Acknowledgements

This project has been made possible by the Brain Canada Foundation, through the Canada Brain Research Fund, with the financial support of Health Canada, National Institutes of Health (NIH R01 AG068563A), and the Canadian Institute of Health Research (CIHR 438531). DB was also supported by the Healthy Brains Healthy Lives initiative (Canada First Research Excellence fund), Google (Research Award & Teaching Award), and by the CIFAR Artificial Intelligence Chairs program (Canada Institute for Advanced Research).

## Tables

**Table 1:** Main effects of solitary living. Shows marginal posterior parameter distributions.

|  | mean | hdi_10% | hdi_90% |
| --- | --- | --- | --- |
| interc[0] | -1.232 | -1.253 | -1.209 |
| interc[1] | -1.69 | -1.717 | -1.665 |
| age[0] | 0.311 | 0.288 | 0.334 |
| age[1] | 0.012 | -0.014 | 0.038 |
| LH_Vis_1[0] | -0.064 | -0.112 | 0.0 |
| LH_Vis_2[0] | 0.002 | -0.043 | 0.044 |
| LH_Vis_3[0] | 0.062 | -0.001 | 0.111 |
| LH_Vis_4[0] | 0.025 | -0.019 | 0.068 |
| LH_Vis_5[0] | -0.009 | -0.049 | 0.03 |
| LH_Vis_6[0] | -0.029 | -0.067 | 0.013 |
| LH_Vis_7[0] | 0.013 | -0.031 | 0.053 |
| LH_Vis_8[0] | 0.012 | -0.034 | 0.055 |
| LH_Vis_9[0] | -0.039 | -0.084 | 0.006 |
| LH_SomMot_1[0] | 0.012 | -0.034 | 0.061 |
| LH_SomMot_2[0] | 0.065 | -0.001 | 0.113 |
| LH_SomMot_3[0] | -0.018 | -0.065 | 0.031 |
| LH_SomMot_4[0] | 0.052 | -0.003 | 0.096 |
| LH_SomMot_5[0] | -0.02 | -0.06 | 0.024 |
| LH_SomMot_6[0] | 0.012 | -0.031 | 0.058 |
| LH_DorsAttn_Post_1[0] | -0.001 | -0.029 | 0.028 |
| LH_DorsAttn_Post_2[0] | -0.01 | -0.036 | 0.017 |
| LH_DorsAttn_Post_3[0] | 0.009 | -0.019 | 0.036 |
| LH_DorsAttn_Post_4[0] | -0.003 | -0.027 | 0.02 |
| LH_DorsAttn_Post_5[0] | -0.013 | -0.037 | 0.013 |
| LH_DorsAttn_Post_6[0] | -0.015 | -0.039 | 0.013 |
| LH_DorsAttn_PrCv_1[0] | -0.009 | -0.037 | 0.018 |
| LH_DorsAttn_FEF_1[0] | -0.016 | -0.044 | 0.015 |
| LH_SalVentAttn_ParOper_1[0] | -0.022 | -0.053 | 0.02 |
| LH_SalVentAttn_FrOperIns_1[0] | -0.018 | -0.051 | 0.025 |
| LH_SalVentAttn_FrOperIns_2[0] | 0.027 | -0.016 | 0.068 |
| LH_SalVentAttn_PFCI_1[0] | 0.012 | -0.026 | 0.044 |
| LH_SalVentAttn_Med_1[0] | -0.007 | -0.046 | 0.026 |
| LH_SalVentAttn_Med_2[0] | 0.01 | -0.02 | 0.041 |
| LH_SalVentAttn_Med_3[0] | 0.008 | -0.024 | 0.04 |
| LH_Limbic_OFC_1[0] | -0.002 | -0.046 | 0.045 |
| LH_Limbic_TempPole_1[0] | -0.043 | -0.094 | 0.016 |
| LH_Limbic_TempPole_2[0] | -0.015 | -0.053 | 0.026 |
| LH_Cont_Par_1[0] | -0.002 | -0.024 | 0.022 |
| LH_Cont_PFC1_1[0] | -0.007 | -0.031 | 0.018 |
| LH_Cont_pCun_1[0] | -0.02 | -0.044 | 0.008 |
| LH_Cont_Cing_1[0] | 0.01 | -0.012 | 0.033 |
| LH_Default_Temp_1[0] | 0.059 | 0.0 | 0.116 |
| LH_Default_Temp_2[0] | -0.022 | -0.073 | 0.03 |
| LH_Default_Par_1[0] | -0.076 | -0.125 | -0.021 |
| LH_Default_Par_2[0] | 0.008 | -0.041 | 0.058 |
| LH_Default_PFC_1[0] | 0.014 | -0.039 | 0.068 |
| LH_Default_PFC_2[0] | -0.01 | -0.067 | 0.044 |
| LH_Default_PFC_3[0] | -0.038 | -0.094 | 0.024 |
| LH_Default_PFC_4[0] | -0.038 | -0.087 | 0.009 |
| LH_Default_PFC_5[0] | 0.0 | -0.048 | 0.052 |
| LH_Default_PFC_6[0] | -0.012 | -0.051 | 0.027 |
| LH_Default_PFC_7[0] | 0.003 | -0.041 | 0.044 |
| LH_Default_pCunPCC_1[0] | -0.037 | -0.083 | 0.008 |
| LH_Default_pCunPCC_2[0] | 0.003 | -0.046 | 0.054 |
| RH_Vis_1[0] | -0.017 | -0.065 | 0.033 |
| RH_Vis_2[0] | 0.033 | -0.023 | 0.079 |
| RH_Vis_3[0] | -0.046 | -0.095 | 0.009 |
| RH_Vis_4[0] | -0.026 | -0.067 | 0.019 |
| RH_Vis_5[0] | -0.026 | -0.068 | 0.019 |
| RH_Vis_6[0] | -0.031 | -0.078 | 0.015 |
| RH_Vis_7[0] | 0.032 | -0.015 | 0.074 |
| RH_Vis_8[0] | 0.007 | -0.031 | 0.049 |
| RH_SomMot_1[0] | 0.028 | -0.024 | 0.077 |
| RH_SomMot_2[0] | -0.022 | -0.068 | 0.029 |
| RH_SomMot_3[0] | -0.04 | -0.086 | 0.008 |
| RH_SomMot_4[0] | 0.014 | -0.026 | 0.055 |
| RH_SomMot_5[0] | -0.048 | -0.088 | 0.003 |
| RH_SomMot_6[0] | -0.013 | -0.051 | 0.027 |
| RH_SomMot_7[0] | -0.033 | -0.067 | 0.005 |
| RH_SomMot_8[0] | 0.031 | -0.015 | 0.074 |
| RH_DorsAttn_Post_1[0] | 0.008 | -0.023 | 0.035 |
| RH_DorsAttn_Post_2[0] | 0.006 | -0.021 | 0.03 |
| RH_DorsAttn_Post_3[0] | 0.004 | -0.021 | 0.029 |
| RH_DorsAttn_Post_4[0] | 0.028 | -0.008 | 0.059 |
| RH_DorsAttn_Post_5[0] | 0.001 | -0.023 | 0.026 |
| RH_DorsAttn_PrCv_1[0] | -0.007 | -0.035 | 0.02 |
| RH_DorsAttn_FEF_1[0] | 0.017 | -0.012 | 0.044 |
| RH_SalVentAttn_TempOccPar_1[0] | 0.044 | -0.007 | 0.087 |
| RH_SalVentAttn_TempOccPar_2[0] | -0.005 | -0.04 | 0.03 |
| RH_SalVentAttn_FrOperIns_1[0] | -0.016 | -0.057 | 0.026 |
| RH_SalVentAttn_Med_1[0] | 0.024 | -0.012 | 0.056 |
| RH_SalVentAttn_Med_2[0] | -0.012 | -0.045 | 0.026 |
| RH_Limbic_OFC_1[0] | 0.014 | -0.034 | 0.058 |
| RH_Limbic_TempPole_1[0] | -0.01 | -0.057 | 0.037 |
| RH_Cont_Par_1[0] | 0.002 | -0.02 | 0.023 |
| RH_Cont_Par_2[0] | -0.002 | -0.023 | 0.02 |
| RH_Cont_PFCI_1[0] | -0.009 | -0.033 | 0.017 |
| RH_Cont_PFCI_2[0] | -0.003 | -0.027 | 0.019 |
| RH_Cont_PFCI_3[0] | 0.001 | -0.023 | 0.023 |
| RH_Cont_PFCI_4[0] | -0.01 | -0.033 | 0.015 |
| RH_Cont_Cing_1[0] | 0.003 | -0.019 | 0.026 |
| RH_Cont_PFCmp_1[0] | -0.004 | -0.027 | 0.021 |
| RH_Cont_pCun_1[0] | -0.001 | -0.025 | 0.021 |
| RH_Default_Par_1[0] | 0.001 | -0.042 | 0.048 |
| RH_Default_Temp_1[0] | 0.089 | 0.033 | 0.147 |
| RH_Default_Temp_2[0] | 0.086 | 0.031 | 0.145 |
| RH_Default_Temp_3[0] | 0.08 | 0.019 | 0.136 |
| RH_Default_PFCv_1[0] | -0.016 | -0.072 | 0.04 |
| RH_Default_PFCv_2[0] | 0.062 | 0.015 | 0.113 |
| RH_Default_PFCdPFCm_1[0] | -0.014 | -0.067 | 0.047 |
| RH_Default_PFCdPFCm_2[0] | 0.029 | -0.023 | 0.076 |
| RH_Default_PFCdPFCm_3[0] | -0.097 | -0.144 | -0.047 |
| RH_Default_pCunPCC_1[0] | -0.036 | -0.086 | 0.008 |
| RH_Default_pCunPCC_2[0] | 0.043 | -0.009 | 0.093 |
| sigma_nets_Cont[0] | 0.021 | 0.001 | 0.031 |
| sigma_nets_Default[0] | 0.065 | 0.044 | 0.083 |
| sigma_nets_DorsAttn[0] | 0.025 | 0.001 | 0.038 |
| sigma_nets_Limbic[0] | 0.054 | 0.001 | 0.081 |
| sigma_nets_SalVentAttn[0] | 0.037 | 0.002 | 0.055 |
| sigma_nets_SomMot[0] | 0.051 | 0.019 | 0.076 |
| sigma_nets_Vis[0] | 0.05 | 0.021 | 0.074 |

**Table 2:** Sex effects of solitary living. Shows marginal posterior parameter distributions (male - female effect).

|  | mean | hdi_25% | hdi_75% |
| --- | --- | --- | --- |
| interc | -0.462 | -0.481 | -0.446 |
| age | -0.3 | -0.316 | -0.28 |
| sigma_nets_Cont_log__ | 0.543 | -0.223 | 1.257 |
| sigma_nets_Default_log__ | 0.142 | -0.214 | 0.442 |
| sigma_nets_DorsAttn_log__ | 0.35 | -0.641 | 0.883 |
| sigma_nets_Limbic_log__ | -0.199 | -1.073 | 0.691 |
| sigma_nets_SalVentAttn_log__ | 0.442 | -0.334 | 0.965 |
| sigma_nets_SomMot_log__ | -0.54 | -1.038 | 0.246 |
| sigma_nets_Vis_log__ | 0.394 | -0.314 | 0.943 |
| LH_Vis_1 | -0.05 | -0.085 | 0.013 |
| LH_Vis_2 | 0.051 | -0.009 | 0.071 |
| LH_Vis_3 | -0.016 | -0.055 | 0.033 |
| LH_Vis_4 | 0.001 | -0.043 | 0.029 |
| LH_Vis_5 | -0.042 | -0.067 | 0.002 |
| LH_Vis_6 | 0.007 | -0.031 | 0.039 |
| LH_Vis_7 | -0.028 | -0.053 | 0.018 |
| LH_Vis_8 | 0.042 | -0.006 | 0.07 |
| LH_Vis_9 | -0.042 | -0.065 | 0.016 |
| LH_SomMot_1 | 0.024 | -0.021 | 0.051 |
| LH_SomMot_2 | -0.028 | -0.059 | 0.024 |
| LH_SomMot_3 | -0.008 | -0.041 | 0.03 |
| LH_SomMot_4 | -0.042 | -0.067 | 0.013 |
| LH_SomMot_5 | 0.05 | 0.003 | 0.075 |
| LH_SomMot_6 | 0.004 | -0.03 | 0.037 |
| LH_DorsAttn_Post_1 | 0.025 | -0.014 | 0.041 |
| LH_DorsAttn_Post_2 | 0.009 | -0.017 | 0.034 |
| LH_DorsAttn_Post_3 | -0.004 | -0.027 | 0.024 |
| LH_DorsAttn_Post_4 | 0.02 | -0.015 | 0.034 |
| LH_DorsAttn_Post_5 | 0.019 | -0.014 | 0.036 |
| LH_DorsAttn_Post_6 | -0.015 | -0.035 | 0.015 |
| LH_DorsAttn_PrCv_1 | 0.011 | -0.02 | 0.031 |
| LH_DorsAttn_FEF_1 | -0.029 | -0.046 | 0.012 |
| LH_SalVentAttn_ParOper_1 | -0.009 | -0.044 | 0.031 |
| LH_SalVentAttn_FrOperIns_1 | -0.067 | -0.083 | 0.006 |
| LH_SalVentAttn_FrOperIns_2 | 0.075 | -0.007 | 0.093 |
| LH_SalVentAttn_PFCI_1 | -0.048 | -0.075 | 0.001 |
| LH_SalVentAttn_Med_1 | 0.008 | -0.031 | 0.041 |
| LH_SalVentAttn_Med_2 | 0.026 | -0.013 | 0.05 |
| LH_SalVentAttn_Med_3 | 0.003 | -0.026 | 0.039 |
| LH_Limbic_OFC_1 | -0.015 | -0.054 | 0.032 |
| LH_Limbic_TempPole_1 | -0.005 | -0.049 | 0.046 |
| LH_Limbic_TempPole_2 | 0.036 | -0.013 | 0.062 |
| LH_Cont_Par_1 | 0.003 | -0.021 | 0.023 |
| LH_Cont_PFC1_1 | 0.023 | -0.014 | 0.036 |
| LH_Cont_pCun_1 | -0.031 | -0.045 | 0.011 |
| LH_Cont_Cing_1 | -0.003 | -0.025 | 0.02 |
| LH_Default_Temp_1 | 0.047 | -0.002 | 0.098 |
| LH_Default_Temp_2 | 0.009 | -0.034 | 0.052 |
| LH_Default_Par_1 | -0.005 | -0.053 | 0.038 |
| LH_Default_Par_2 | -0.023 | -0.064 | 0.018 |
| LH_Default_PFC_1 | 0.017 | -0.023 | 0.067 |
| LH_Default_PFC_2 | -0.004 | -0.053 | 0.038 |
| LH_Default_PFC_3 | 0.018 | -0.03 | 0.065 |
| LH_Default_PFC_4 | -0.07 | -0.109 | -0.02 |
| LH_Default_PFC_5 | -0.01 | -0.051 | 0.033 |
| LH_Default_PFC_6 | -0.093 | -0.127 | -0.05 |
| LH_Default_PFC_7 | -0.032 | -0.068 | 0.009 |
| LH_Default_pCunPCC_1 | 0.032 | -0.01 | 0.072 |
| LH_Default_pCunPCC_2 | -0.048 | -0.086 | 0.002 |
| RH_Vis_1 | 0.04 | -0.016 | 0.066 |
| RH_Vis_2 | -0.037 | -0.063 | 0.019 |
| RH_Vis_3 | 0.011 | -0.03 | 0.052 |
| RH_Vis_4 | -0.051 | -0.074 | 0.007 |
| RH_Vis_5 | -0.021 | -0.058 | 0.017 |
| RH_Vis_6 | 0.007 | -0.031 | 0.043 |
| RH_Vis_7 | -0.008 | -0.041 | 0.034 |
| RH_Vis_8 | 0.075 | 0.016 | 0.099 |
| RH_SomMot_1 | -0.011 | -0.047 | 0.029 |
| RH_SomMot_2 | 0.076 | 0.013 | 0.103 |
| RH_SomMot_3 | 0.012 | -0.025 | 0.045 |
| RH_SomMot_4 | -0.007 | -0.04 | 0.024 |
| RH_SomMot_5 | 0.055 | 0.006 | 0.083 |
| RH_SomMot_6 | 0.027 | -0.008 | 0.054 |
| RH_SomMot_7 | -0.013 | -0.035 | 0.026 |
| RH_SomMot_8 | -0.005 | -0.04 | 0.029 |
| RH_DorsAttn_Post_1 | -0.019 | -0.038 | 0.018 |
| RH_DorsAttn_Post_2 | 0.017 | -0.014 | 0.035 |
| RH_DorsAttn_Post_3 | -0.019 | -0.04 | 0.009 |
| RH_DorsAttn_Post_4 | 0.042 | -0.014 | 0.053 |
| RH_DorsAttn_Post_5 | -0.008 | -0.026 | 0.021 |
| RH_DorsAttn_PrCv_1 | 0.015 | -0.015 | 0.037 |
| RH_DorsAttn_FEF_1 | -0.051 | -0.061 | 0.003 |
| RH_SalVentAttn_TempOccPar_1 | -0.048 | -0.081 | 0.012 |
| RH_SalVentAttn_TempOccPar_2 | -0.028 | -0.053 | 0.019 |
| RH_SalVentAttn_FrOperIns_1 | -0.026 | -0.059 | 0.023 |
| RH_SalVentAttn_Med_1 | 0.052 | 0.004 | 0.079 |
| RH_SalVentAttn_Med_2 | 0.025 | -0.023 | 0.049 |
| RH_Limbic_OFC_1 | -0.065 | -0.081 | 0.013 |
| RH_Limbic_TempPole_1 | 0.026 | -0.029 | 0.057 |
| RH_Cont_Par_1 | -0.004 | -0.026 | 0.017 |
| RH_Cont_Par_2 | -0.019 | -0.034 | 0.011 |
| RH_Cont_PFCI_1 | -0.019 | -0.032 | 0.019 |
| RH_Cont_PFCI_2 | 0.014 | -0.016 | 0.03 |
| RH_Cont_PFCI_3 | 0.008 | -0.018 | 0.028 |
| RH_Cont_PFCI_4 | -0.03 | -0.042 | 0.011 |
| RH_Cont_Cing_1 | -0.009 | -0.028 | 0.017 |
| RH_Cont_PFCmp_1 | 0.005 | -0.019 | 0.027 |
| RH_Cont_pCun_1 | 0.019 | -0.014 | 0.033 |
| RH_Default_Par_1 | -0.04 | -0.076 | 0.004 |
| RH_Default_Temp_1 | -0.021 | -0.066 | 0.031 |
| RH_Default_Temp_2 | 0.038 | -0.014 | 0.085 |
| RH_Default_Temp_3 | -0.014 | -0.059 | 0.041 |
| RH_Default_PFCv_1 | -0.001 | -0.048 | 0.042 |
| RH_Default_PFCv_2 | 0.013 | -0.033 | 0.054 |
| RH_Default_PFCdPFCm_1 | 0.04 | -0.017 | 0.078 |
| RH_Default_PFCdPFCm_2 | -0.01 | -0.053 | 0.03 |
| RH_Default_PFCdPFCm_3 | 0.024 | -0.023 | 0.068 |
| RH_Default_pCunPCC_1 | 0.046 | 0.0 | 0.083 |
| RH_Default_pCunPCC_2 | 0.059 | 0.008 | 0.099 |
| sigma_nets_Cont | 0.015 | -0.005 | 0.031 |
| sigma_nets_Default | 0.009 | -0.011 | 0.028 |
| sigma_nets_DorsAttn | 0.009 | -0.017 | 0.024 |
| sigma_nets_Limbic | -0.009 | -0.055 | 0.03 |
| sigma_nets_SalVentAttn | 0.019 | -0.009 | 0.047 |
| sigma_nets_SomMot | -0.021 | -0.046 | 0.005 |
| sigma_nets_Vis | 0.017 | -0.009 | 0.045 |

**Table 3:** Main effects of solitary living in amygdala nuclei. Shows marginal posterior parameter distributions.

|  | mean | hdi_10% | hdi_90% |
| --- | --- | --- | --- |
| interc[0] | -1.229 | -1.25 | -1.207 |
| interc[1] | -1.691 | -1.717 | -1.665 |
| age[0] | 0.313 | 0.29 | 0.335 |
| age[1] | 0.008 | -0.018 | 0.034 |
| Volume of Accessory-Basal-nucleus (left hemisphere)[0] | -0.109 | -0.196 | -0.018 |
| Volume of Accessory-Basal-nucleus (right hemisphere)[0] | 0.224 | 0.136 | 0.319 |
| Volume of Anterior-amygdaloid-area-AAA (left hemisphere)[0] | -0.006 | -0.037 | 0.024 |
| Volume of Anterior-amygdaloid-area-AAA (right hemisphere)[0] | 0.018 | -0.013 | 0.05 |
| Volume of Basal-nucleus (left hemisphere)[0] | 0.021 | -0.068 | 0.12 |
| Volume of Basal-nucleus (right hemisphere)[0] | -0.063 | -0.15 | 0.033 |
| Volume of Central-nucleus (left hemisphere)[0] | 0.018 | -0.014 | 0.046 |
| Volume of Central-nucleus (right hemisphere)[0] | -0.048 | -0.078 | -0.017 |
| Volume of Cortical-nucleus (left hemisphere)[0] | 0.039 | 0.0 | 0.078 |
| Volume of Cortical-nucleus (right hemisphere)[0] | -0.057 | -0.095 | -0.019 |
| Volume of Corticoamygdaloid-transitio (left hemisphere)[0] | 0.037 | -0.004 | 0.083 |
| Volume of Corticoamygdaloid-transitio (right hemisphere)[0] | -0.119 | -0.167 | -0.072 |
| Volume of Lateral-nucleus (left hemisphere)[0] | 0.0 | -0.048 | 0.045 |
| Volume of Lateral-nucleus (right hemisphere)[0] | 0.03 | -0.016 | 0.075 |
| Volume of Medial-nucleus (left hemisphere)[0] | -0.022 | -0.051 | 0.007 |
| Volume of Medial-nucleus (right hemisphere)[0] | 0.007 | -0.022 | 0.038 |
| Volume of Paralaminar-nucleus (left hemisphere)[0] | -0.007 | -0.055 | 0.042 |
| Volume of Paralaminar-nucleus (right hemisphere)[0] | 0.006 | -0.043 | 0.052 |

**Table 4:**
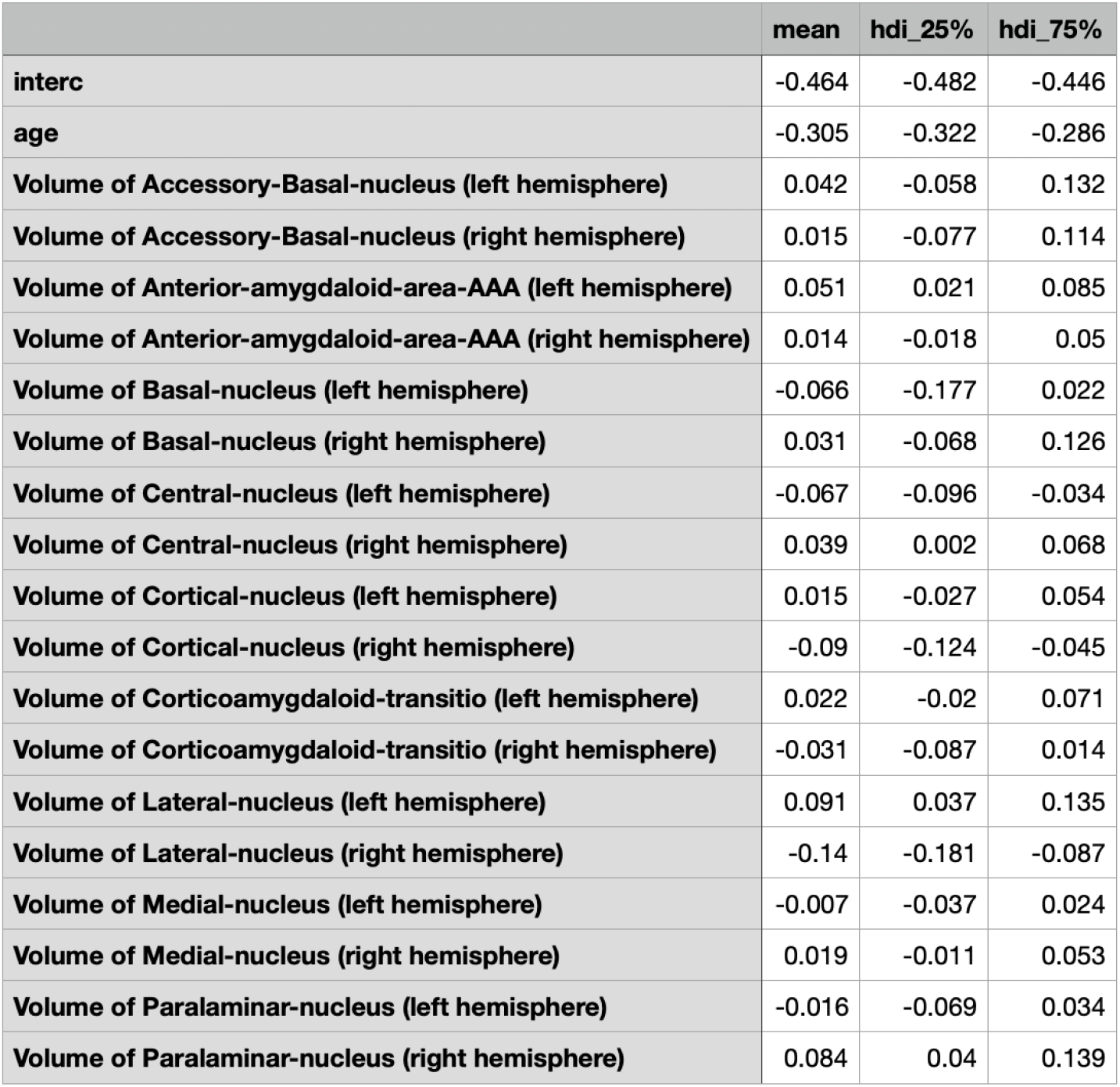
Sex differentiation of solitary living in amygdala nuclei. Shows marginal posterior parameter distributions (male - female effect).

|  | mean | hdi_25% | hdi_75% |
| --- | --- | --- | --- |
| interc | -0.464 | -0.482 | -0.446 |
| age | -0.305 | -0.322 | -0.286 |
| Volume of Accessory-Basal-nucleus (left hemisphere) | 0.042 | -0.058 | 0.132 |
| Volume of Accessory-Basal-nucleus (right hemisphere) | 0.015 | -0.077 | 0.114 |
| Volume of Anterior-amygdaloid-area-AAA (left hemisphere) | 0.051 | 0.021 | 0.085 |
| Volume of Anterior-amygdaloid-area-AAA (right hemisphere) | 0.014 | -0.018 | 0.05 |
| Volume of Basal-nucleus (left hemisphere) | -0.066 | -0.177 | 0.022 |
| Volume of Basal-nucleus (right hemisphere) | 0.031 | -0.068 | 0.126 |
| Volume of Central-nucleus (left hemisphere) | -0.067 | -0.096 | -0.034 |
| Volume of Central-nucleus (right hemisphere) | 0.039 | 0.002 | 0.068 |
| Volume of Cortical-nucleus (left hemisphere) | 0.015 | -0.027 | 0.054 |
| Volume of Cortical-nucleus (right hemisphere) | -0.09 | -0.124 | -0.045 |
| Volume of Corticoamygdaloid-transitio (left hemisphere) | 0.022 | -0.02 | 0.071 |
| Volume of Corticoamygdaloid-transitio (right hemisphere) | -0.031 | -0.087 | 0.014 |
| Volume of Lateral-nucleus (left hemisphere) | 0.091 | 0.037 | 0.135 |
| Volume of Lateral-nucleus (right hemisphere) | -0.14 | -0.181 | -0.087 |
| Volume of Medial-nucleus (left hemisphere) | -0.007 | -0.037 | 0.024 |
| Volume of Medial-nucleus (right hemisphere) | 0.019 | -0.011 | 0.053 |
| Volume of Paralaminar-nucleus (left hemisphere) | -0.016 | -0.069 | 0.034 |
| Volume of Paralaminar-nucleus (right hemisphere) | 0.084 | 0.04 | 0.139 |

**Table 5:** Main effects of solitary living in hippocampus nuclei groups. Shows marginal posterior parameter distributions.

|  | mean | hdi_10% | hdi_90% |
| --- | --- | --- | --- |
| interc[0] | -1.232 | -1.254 | -1.211 |
| interc[1] | -1.696 | -1.721 | -1.669 |
| age[0] | 0.313 | 0.29 | 0.336 |
| age[1] | 0.008 | -0.016 | 0.036 |
| Volume of CA1-body (left hemisphere)[0] | -0.03 | -0.063 | 0.002 |
| Volume of CA1-body (right hemisphere)[0] | -0.02 | -0.052 | 0.012 |
| Volume of CA1-head (left hemisphere)[0] | 0.048 | 0.003 | 0.094 |
| Volume of CA1-head (right hemisphere)[0] | 0.003 | -0.045 | 0.049 |
| Volume of CA3-body (left hemisphere)[0] | -0.031 | -0.066 | 0.006 |
| Volume of CA3-body (right hemisphere)[0] | 0.027 | -0.007 | 0.062 |
| Volume of CA3-head (left hemisphere)[0] | 0.016 | -0.025 | 0.051 |
| Volume of CA3-head (right hemisphere)[0] | -0.09 | -0.127 | -0.054 |
| Volume of CA4-body (left hemisphere)[0] | 0.094 | 0.018 | 0.175 |
| Volume of CA4-body (right hemisphere)[0] | 0.07 | -0.014 | 0.152 |
| Volume of CA4-head (left hemisphere)[0] | 0.077 | -0.021 | 0.176 |
| Volume of CA4-head (right hemisphere)[0] | -0.015 | -0.11 | 0.082 |
| Volume of HATA (left hemisphere)[0] | 0.003 | -0.02 | 0.027 |
| Volume of HATA (right hemisphere)[0] | 0.004 | -0.019 | 0.027 |
| Volume of Hippocampal-tail (left hemisphere)[0] | -0.046 | -0.076 | -0.015 |
| Volume of Hippocampal-tail (right hemisphere)[0] | 0.046 | 0.016 | 0.079 |
| Volume of fimbria (left hemisphere)[0] | -0.02 | -0.045 | 0.004 |
| Volume of fimbria (right hemisphere)[0] | -0.036 | -0.059 | -0.014 |
| Volume of parasubiculum (left hemisphere)[0] | -0.005 | -0.03 | 0.018 |
| Volume of parasubiculum (right hemisphere)[0] | -0.009 | -0.033 | 0.016 |
| Volume of presubiculum-body (left hemisphere)[0] | -0.044 | -0.079 | -0.01 |
| Volume of presubiculum-body (right hemisphere)[0] | 0.046 | 0.013 | 0.08 |
| Volume of presubiculum-head (left hemisphere)[0] | 0.054 | 0.014 | 0.096 |
| Volume of presubiculum-head (right hemisphere)[0] | 0.043 | 0.002 | 0.084 |
| Volume of subiculum-body (left hemisphere)[0] | -0.009 | -0.045 | 0.029 |
| Volume of subiculum-body (right hemisphere)[0] | 0.01 | -0.028 | 0.048 |
| Volume of subiculum-head (left hemisphere)[0] | -0.036 | -0.077 | 0.008 |
| Volume of subiculum-head (right hemisphere)[0] | -0.041 | -0.087 | 0.001 |
| Volume of hippocampal-fissure (left hemisphere)[0] | 0.019 | -0.008 | 0.047 |
| Volume of hippocampal-fissure (right hemisphere)[0] | -0.016 | -0.045 | 0.011 |
| Volume of molecular-layer-HP-body (left hemisphere)[0] | 0.017 | -0.011 | 0.046 |
| Volume of molecular-layer-HP-body (right hemisphere)[0] | -0.013 | -0.041 | 0.015 |
| Volume of molecular-layer-HP-head (left hemisphere)[0] | -0.034 | -0.064 | -0.003 |
| Volume of molecular-layer-HP-head (right hemisphere)[0] | 0.04 | 0.008 | 0.071 |
| Volume of GC-ML-DG-body (left hemisphere)[0] | -0.048 | -0.119 | 0.024 |
| Volume of GC-ML-DG-body (right hemisphere)[0] | -0.113 | -0.192 | -0.038 |
| Volume of GC-ML-DG-head (left hemisphere)[0] | -0.152 | -0.256 | -0.046 |
| Volume of GC-ML-DG-head (right hemisphere)[0] | 0.099 | 0.0 | 0.203 |

**Table 6:** Sex differentiation of solitary living in hippocampus nuclei groups. Shows marginal posterior parameter distributions (male - female effect).

|  | mean | hdi_25% | hdi_75% |
| --- | --- | --- | --- |
| interc | -0.469 | -0.487 | -0.451 |
| age | -0.305 | -0.325 | -0.288 |
| Volume of CA1-body (left hemisphere) | 0.001 | -0.033 | 0.037 |
| Volume of CA1-body (right hemisphere) | -0.016 | -0.05 | 0.018 |
| Volume of CA1-head (left hemisphere) | 0.032 | -0.012 | 0.083 |
| Volume of CA1-head (right hemisphere) | -0.018 | -0.069 | 0.029 |
| Volume of CA3-body (left hemisphere) | -0.023 | -0.059 | 0.015 |
| Volume of CA3-body (right hemisphere) | -0.008 | -0.048 | 0.027 |
| Volume of CA3-head (left hemisphere) | 0.042 | -0.003 | 0.077 |
| Volume of CA3-head (right hemisphere) | -0.117 | -0.157 | -0.08 |
| Volume of CA4-body (left hemisphere) | -0.012 | -0.1 | 0.063 |
| Volume of CA4-body (right hemisphere) | 0.133 | 0.036 | 0.215 |
| Volume of CA4-head (left hemisphere) | -0.12 | -0.23 | -0.018 |
| Volume of CA4-head (right hemisphere) | -0.079 | -0.192 | 0.012 |
| Volume of HATA (left hemisphere) | -0.015 | -0.044 | 0.007 |
| Volume of HATA (right hemisphere) | 0.001 | -0.023 | 0.028 |
| Volume of Hippocampal-tail (left hemisphere) | 0.027 | -0.011 | 0.053 |
| Volume of Hippocampal-tail (right hemisphere) | -0.066 | -0.098 | -0.034 |
| Volume of fimbria (left hemisphere) | 0.043 | 0.017 | 0.069 |
| Volume of fimbria (right hemisphere) | -0.001 | -0.024 | 0.024 |
| Volume of parasubiculum (left hemisphere) | -0.027 | -0.055 | -0.003 |
| Volume of parasubiculum (right hemisphere) | 0.1 | 0.076 | 0.128 |
| Volume of presubiculum-body (left hemisphere) | -0.054 | -0.095 | -0.022 |
| Volume of presubiculum-body (right hemisphere) | 0.032 | -0.002 | 0.07 |
| Volume of presubiculum-head (left hemisphere) | -0.034 | -0.083 | 0.005 |
| Volume of presubiculum-head (right hemisphere) | -0.014 | -0.057 | 0.03 |
| Volume of subiculum-body (left hemisphere) | -0.007 | -0.048 | 0.029 |
| Volume of subiculum-body (right hemisphere) | -0.086 | -0.127 | -0.048 |
| Volume of subiculum-head (left hemisphere) | 0.03 | -0.009 | 0.081 |
| Volume of subiculum-head (right hemisphere) | -0.011 | -0.06 | 0.034 |
| Volume of hippocampal-fissure (left hemisphere) | -0.012 | -0.04 | 0.018 |
| Volume of hippocampal-fissure (right hemisphere) | 0.063 | 0.032 | 0.092 |
| Volume of molecular-layer-HP-body (left hemisphere) | 0.113 | 0.08 | 0.14 |
| Volume of molecular-layer-HP-body (right hemisphere) | -0.047 | -0.076 | -0.017 |
| Volume of molecular-layer-HP-head (left hemisphere) | 0.028 | -0.005 | 0.063 |
| Volume of molecular-layer-HP-head (right hemisphere) | 0.039 | 0.006 | 0.074 |
| Volume of GC-ML-DG-body (left hemisphere) | 0.054 | -0.021 | 0.131 |
| Volume of GC-ML-DG-body (right hemisphere) | -0.106 | -0.185 | -0.024 |
| Volume of GC-ML-DG-head (left hemisphere) | 0.042 | -0.077 | 0.147 |
| Volume of GC-ML-DG-head (right hemisphere) | 0.137 | 0.029 | 0.241 |

## Supplementary Figures

**Figure S1.**
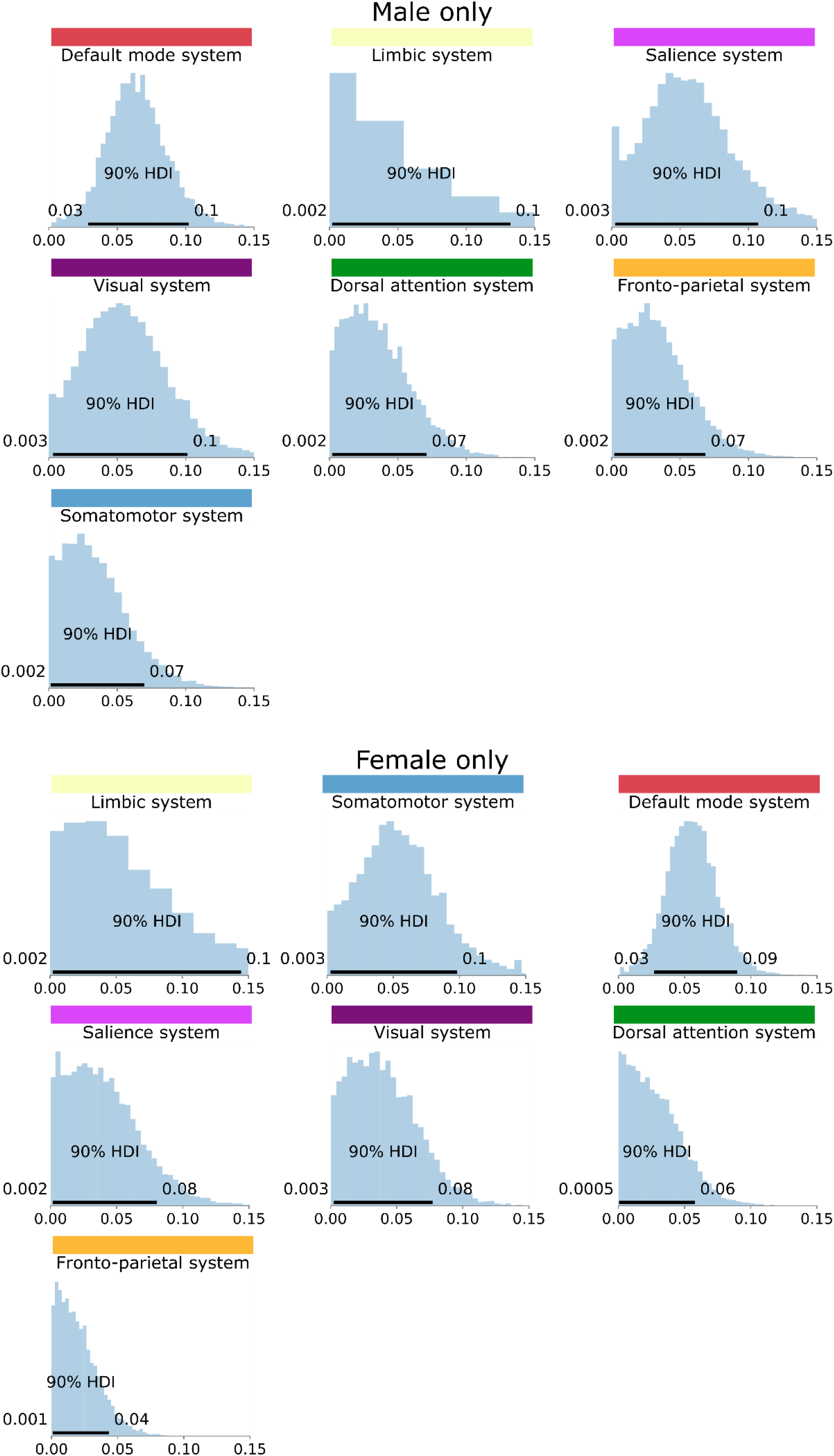
Network level Sex differences: The Bayesian hierarchical modelling framework estimated the gray matter effects for distributed networks of brain regions in explaining living alone in male (top) and female (bottom) subgroups. Histograms show the inferred marginal posterior parameter distributions of the overall explanatory variance (sigma parameter) for each major brain network (volume measures in standard units). Horizontal black bars indicate the highest posterior density interval (HPI) of the model’s network variance parameters, ranging from 10 to 90% probability. Posterior distributions for the variance parameter (sigma) of each brain network are ordered from strongest (top left) to weakest (bottom right) for each sex.

## Notes

### Competing Interest Statement

The authors have declared no competing interest.

## References

Adolphs, R. (2009) The social brain: neural basis of social knowledge. Annu Rev Psychol, 60, 693–716.

Aggleton, J.P., Burton, M.J. & Passingham, R.E. (1980) Cortical and subcortical afferents to the amygdala of the rhesus monkey (Macaca mulatta). Brain Res, 190, 347–368.

Alfaro-Almagro, F., Jenkinson, M., Bangerter, N.K., Andersson, J.L.R., Griffanti, L., Douaud, G., Sotiropoulos, S.N., Jbabdi, S., Hernandez-Fernandez, M., Vallee, E., Vidaurre, D., Webster, M., McCarthy, P., Rorden, C., Daducci, A., Alexander, D.C., Zhang, H., Dragonu, I., Matthews, P.M., Miller, K.L. & Smith, S.M. (2018) Image processing and Quality Control for the first 10,000 brain imaging datasets from UK Biobank. Neuroimage, 166, 400–424.

Amici, F., Kulik, L., Langos, D. & Widdig, A. (2019) Growing into adulthood - a review on sex differences in the development of scoiality across macaques. Behavioural Ecology and Sociobiology, 73.

Amici, F. & Widdig, A. (2019) An evolutionary perspective on the development of primate sociality. Springer.

Anacker, C., Luna, V.M., Stevens, G.S., Millette, A., Shores, R., Jimenez, J.C., Chen, B. & Hen, R. (2018) Hippocampal neurogenesis confers stress resilience by inhibiting the ventral dentate gyrus. Nature, 559, 98–102.

Anderson, J.S., Nielsen, J.A., Froehlich, A.L., DuBray, M.B., Druzgal, T.J., Cariello, A.N., Cooperrider, J.R., Zielinski, B.A., Ravichandran, C., Fletcher, P.T., Alexander, A.L., Bigler, E.D., Lange, N. & Lainhart, J.E. (2011) Functional connectivity magnetic resonance imaging classification of autism. Brain, 134, 3742–3754.

Andersson, J., Jenkinson, M. & Smith, S. (2007) Non-linear optimisation FMRIB Technical Report TR07JA1.

Asperholm, M., Högman, N., Rafi, J. & Herlitz, A. (2019) What did you do yesterday? A meta-analysis of sex differences in episodic memory. Psychol Bull, 145, 785–821.

Baron-Cohen, S., O’Riordan, M., Stone, V., Jones, R. & Plaisted, K. (1999) Recognition of faux pas by normally developing children and children with Asperger syndrome or high-functioning autism. J Autism Dev Disord, 29, 407–418.

Behrens, T.E., Hunt, L.T., Woolrich, M.W. & Rushworth, M.F. (2008) Associative learning of social value. Nature, 456, 245–249.

Bhattacharya, K., Ghosh, A., Monsivais, D., Dunbar, R.I. & Kaski, K. (2016) Sex differences in social focus across the life cycle in humans. R Soc Open Sci, 3, 160097.

Bickart, K.C., Wright, C.I., Dautoff, R.J., Dickerson, B.C. & Barrett, L.F. (2011) Amygdala volume and social network size in humans. Nat Neurosci, 14, 163–164.

Biggio, F., Mostallino, M., Talani, G., Locci, V., Mostallino, R., Calandra, G., Sanna, E. & Biggio, G. (2019a) Social enrichment reverses the isolation-induced deficits of neuronal plasticity in the hippocampus of male rats. Neuropharmacology, 151, 45–54.

Biggio, F., Mostallino, M.C., Talani, G., Locci, V., Mostallino, R., Calandra, G., Sanna, E. & Biggio, G. (2019b) Social enrichment reverses the isolation-induced deficits of neuronal plasticity in the hippocampus of male rats. Neuropharmacology, 151, 45–54.

Boccara, C.N., Sargolini, F., Thoresen, V.H., Solstad, T., Witter, M.P., Moser, E.I. & Moser, M.B. (2010) Grid cells in pre- and parasubiculum. Nat Neurosci, 13, 987–994.

Boorman, E.D., Behrens, T.E., Woolrich, M.W. & Rushworth, M.F. (2009) How green is the grass on the other side? Frontopolar cortex and the evidence in favor of alternative courses of action. Neuron, 62, 733–743.

Byron, E. (2019) More Americans Are Living Solo, and Companies Want Their Business.

Bzdok, D. (2017) Classical statistics and statistical learning in imaging neuroscience. Frontiers in neuroscience, 11, 543.

Bzdok, D. & Dunbar, R.I. (2020a) The neurobiology of social distance. Trends in cognitive sciences

Bzdok, D. & Dunbar, R.I.M. (2020b) The Neurobiology of Social Distance. Trends Cogn Sci, 24, 717–733.

Bzdok, D., Eickenberg, M., Varoquaux, G. & Thirion, B. (Year) Hierarchical region-network sparsity for high-dimensional inference in brain imaging. International Conference on Information Processing in Medical Imaging. Springer, City. p. 323–335.

Bzdok, D., Krzywinski, M. & Altman, N. (2017b) Machine learning: a primer. Nature methods, 14, 1119.

Cacioppo, J.T. & Hawkley, L.C. (2009) Perceived social isolation and cognition. Trends Cogn Sci, 13, 447–454.

Cacioppo, J.T., Norris, C.J., Decety, J., Monteleone, G. & Nusbaum, H. (2009) In the eye of the beholder: individual differences in perceived social isolation predict regional brain activation to social stimuli. Journal of cognitive neuroscience, 21, 83–92.

Cloud, D.H., Drucker, E., Browne, A. & Parsons, J. (2015) Public Health and Solitary Confinement in the United States. Am J Public Health, 105, 18–26.

Cyranowski, J.M., Zill, N., Bode, R., Butt, Z., Kelly, M.A., Pilkonis, P.A., Salsman, J.M. & Cella, D. (2013) Assessing social support, companionship, and distress: National Institute of Health (NIH) Toolbox Adult Social Relationship Scales. Health Psychol, 32, 293–301.

Dollinger, S.J. & Malmquist, D. (2009) Reliability and validity of single-item self-reports: with special relevance to college students’ alcohol use, religiosity, study, and social life. J Gen Psychol, 136, 231–241.

Dranovsky, A. & Leonardo, E.D. (2012) Is there a role for young hippocampal neurons in adaptation to stress? Behav Brain Res, 227, 371–375.

Dunbar, R.I. (2016) Do online social media cut through the constraints that limit the size of offline social networks? R Soc Open Sci, 3, 150292.

Dunbar, R.I. & Shultz, S. (2007a) Evolution in the social brain. Science, 317, 1344–1347.

Dunbar, R.I. & Shultz, S. (2007b) Understanding primate brain evolution. Philos Trans R Soc Lond B Biol Sci, 362, 649–658.

Dunbar, R.I.M. (1992) Neocortex Size as a Constraint on Group-Size in Primates. Journal of Human Evolution, 22, 469–493.

Dunbar, R.I.M. (2018) The Anatomy of Friendship. Trends Cogn Sci, 22, 32–51.

Dunbar, R.I.M. & Shultz, S. (2017) Why are there so many explanations for primate brain evolution? Philos Trans R Soc Lond B Biol Sci, 372.

Efron, B. & Tibshirani, R.J. (1994) An Introduction to the bootstrap. Chapman and Hall/CRC.

Eisenberger, N.I., Lieberman, M.D. & Williams, K.D. (2003) Does rejection hurt? An FMRI study of social exclusion. Science, 302, 290–292.

Esteve, A., Reher, D.S., Trevino, R., Zueras, P. & Turu, A. (2020) Living alone over the life course: Cross-National Variations on an emerging issue. Population and Development Review, 46, 20.

Fanselow, M.S. & Dong, H.W. (2010) Are the dorsal and ventral hippocampus functionally distinct structures? Neuron, 65, 7–19.

Frith, C.D. & Frith, U. (2006) The neural basis of mentalizing. Neuron, 50, 531–534.

Gelman, A., Carlin, J.B., Stern, H.S. & Rubin, D.B. (2014) Bayesian Data Analysis. Chapman & Hall/CRC.

Glasser, M.F., Coalson, T.S., Robinson, E.C., Hacker, C.D., Harwell, J., Yacoub, E., Ugurbil, K., Andersson, J., Beckmann, C.F., Jenkinson, M., Smith, S.M. & Van Essen, D.C. (2016) A multi-modal parcellation of human cerebral cortex. Nature, 536, 171–178.

Gould, E., Tanapat, P., McEwen, B.S., Flügge, G. & Fuchs, E. (1998) Proliferation of granule cell precursors in the dentate gyrus of adult monkeys is diminished by stress. Proc Natl Acad Sci U S A, 95, 3168–3171.

Gould, E., Woolley, C.S. & McEwen, B.S. (1990) Short-term glucocorticoid manipulations affect neuronal morphology and survival in the adult dentate gyrus. Neuroscience, 37, 367–375.

Hafkemeijer, A., van der Grond, J. & Rombouts, S.A. (2012) Imaging the default mode network in aging and dementia. Biochim Biophys Acta, 1822, 431–441.

Hawkley, L.C., Browne, M.W. & Cacioppo, J.T. (2005) How can I connect with thee? Let me count the ways. Psychol Sci, 16, 798–804.

Hawkley, L.C., Burleson, M.H., Berntson, G.G. & Cacioppo, J.T. (2003) Loneliness in everyday life: cardiovascular activity, psychosocial context, and health behaviors. J Pers Soc Psychol, 85, 105–120.

Ibi, D., Takuma, K., Koike, H., Mizoguchi, H., Tsuritani, K., Kuwahara, Y., Kamei, H., Nagai, T., Yoneda, Y., Nabeshima, T. & Yamada, K. (2008) Social isolation rearing-induced impairment of the hippocampal neurogenesis is associated with deficits in spatial memory and emotion-related behaviors in juvenile mice. J Neurochem, 105, 921–932.

Iwai, E. & Yukie, M. (1987) Amygdalofugal and amygdalopetal connections with modality-specific visual cortical areas in macaques (Macaca fuscata, M. mulatta, and M. fascicularis). J Comp Neurol, 261, 362–387.

Jackie Tang, N.G., Johnny Truong (2019) Living alone in Canada.

Jackson, S.E., Beard, E., Angus, C., Field, M. & Brown, J. (2021) Moderators of changes in smoking, drinking and quitting behaviour associated with the first COVID-19 lockdown in England. Addiction.

Jacubowski, A., Abeln, V., Vogt, T., Yi, B., Choukèr, A., Fomina, E., Strüder, H.K. & Schneider, S. (2015) The impact of long-term confinement and exercise on central and peripheral stress markers. Physiol Behav, 152, 106–111.

Jenkinson, M., Bannister, P., Brady, M. & Smith, S. (2002) Improved optimization for the robust and accurate linear registration and motion correction of brain images. Neuroimage, 17, 825–841.

Jenkinson, M. & Smith, S. (2001) A global optimisation method for robust affine registration of brain images. Med Image Anal, 5, 143–156.

Kanai, R., Bahrami, B., Roylance, R. & Rees, G. (2012) Online social network size is reflected in human brain structure. Proc Biol Sci, 279, 1327–1334.

Kempermann, G., Kuhn, H.G. & Gage, F.H. (1998) Experience-induced neurogenesis in the senescent dentate gyrus. J Neurosci, 18, 3206–3212.

Key, C. & Ross, C. (1999) Sex differences in energy expenditure in non-human primates. Proc Biol Sci, 266, 2479–2485.

Kiesow, H., Dunbar, R.I.M., Kable, J.W., Kalenscher, T., Vogeley, K., Schilbach, L., Marquand, A.F., Wiecki, T.V. & Bzdok, D. (2020) 10,000 social brains: Sex differentiation in human brain anatomy. Sci Adv, 6, eaaz1170.

Kiesow, H., Uddin, L.Q., Bernhardt, B.C., Kable, J. & Bzdok, D. (2021) Dissecting the midlife crisis: disentangling social, personality and demographic determinants in social brain anatomy. Commun Biol, 4, 728.

Lenroot, R.K. & Giedd, J.N. (2010) Sex differences in the adolescent brain. Brain Cogn, 72, 46–55.

Lewin, C. & Herlitz, A. (2002) Sex differences in face recognition--women’s faces make the difference. Brain Cogn, 50, 121–128.

Lewis, P.A., Rezaie, R., Brown, R., Roberts, N. & Dunbar, R.I. (2011) Ventromedial prefrontal volume predicts understanding of others and social network size. Neuroimage, 57, 1624–1629.

Li, S., Jin, M., Zhang, D., Yang, T., Koeglsperger, T., Fu, H. & Selkoe, D.J. (2013) Environmental novelty activates β2-adrenergic signaling to prevent the impairment of hippocampal LTP by Aβ oligomers. Neuron, 77, 929–941.

Luhmann, M. & Hawkley, L.C. (2016) Age differences in loneliness from late adolescence to oldest old age. Dev Psychol, 52, 943–959.

Magarin, A. & McEwen, B.J.N. (1995) Stress-induced atrophy of apical dendrites of hippocampal CA3c neurons: comparison of stressors. 69, 83–88.

Magariños, A.M., Verdugo, J.M. & McEwen, B.S. (1997a) Chronic stress alters synaptic terminal structure in hippocampus. Proc Natl Acad Sci U S A, 94, 14002–14008.

Magariños, A.M., Verdugo, J.M.G. & McEwen, B.S.J.P.o.t.N.A.o.S. (1997b) Chronic stress alters synaptic terminal structure in hippocampus. 94, 14002–14008.

Magariños, A.M.a., McEwen, B.S., Flügge, G. & Fuchs, E.J.J.o.N. (1996) Chronic psychosocial stress causes apical dendritic atrophy of hippocampal CA3 pyramidal neurons in subordinate tree shrews. 16, 3534–3540.

Mars, R.B., Neubert, F.X., Noonan, M.P., Sallet, J., Toni, I. & Rushworth, M.F. (2012) On the relationship between the “default mode network” and the “social brain”. Front Hum Neurosci, 6, 189.

Mars, R.B., Sallet, J., Neubert, F.X. & Rushworth, M.F. (2013) Connectivity profiles reveal the relationship between brain areas for social cognition in human and monkey temporoparietal cortex. Proc Natl Acad Sci U S A, 110, 10806–10811.

Mashek, D., Cannaday, L.W. & Tangney, J.P. (2007) Inclusion of community in self scale: A single-item pictorial measure of community connectedness. Journal of Community Psychology, 35, 257–275.

Miller, K.L., Alfaro-Almagro, F., Bangerter, N.K., Thomas, D.L., Yacoub, E., Xu, J., Bartsch, A.J., Jbabdi, S., Sotiropoulos, S.N., Andersson, J.L., Griffanti, L., Douaud, G., Okell, T.W., Weale, P., Dragonu, I., Garratt, S., Hudson, S., Collins, R., Jenkinson, M., Matthews, P.M. & Smith, S.M. (2016) Multimodal population brain imaging in the UK Biobank prospective epidemiological study. Nat Neurosci, 19, 1523–1536.

Mishra, M.V., Likitlersuang, J., B Wilmer, J., Cohan, S., Germine, L. & DeGutis, J.M. (2019) Gender Differences in Familiar Face Recognition and the Influence of Sociocultural Gender Inequality. Sci Rep, 9, 17884.

Noonan, M.P., Mars, R.B., Neubert, F.X., Ahmed, B., Smith, J., Krug, K. & Sallet, J. (2016) Organization of the social brain in macaques and humans. In Dreher, J.C., Tremblay, L. (eds) Decision Neuroscience: Handbook of Reward and Decision Making. Elsevier, San Diego.

Noonan, M.P., Mars, R.B., Sallet, J., Dunbar, R.I.M. & Fellows, L.K. (2018) The structural and functional brain networks that support human social networks. Behav Brain Res, 355, 12–23.

OfCom (2020) Media Nations 2020: Interactive Report.

Olderbak, S., Wilhelm, O., Hildebrandt, A. & Quoidbach, J. (2019) Sex differences in facial emotion perception ability across the lifespan. Cogn Emot, 33, 579–588.

Pessoa, L. & Adolphs, R. (2010) Emotion processing and the amygdala: from a ‘low road’ to ‘many roads’ of evaluating biological significance. Nat Rev Neurosci, 11, 773–783.

Proverbio, A.M. (2017) Sex differences in social cognition: The case of face processing. J Neurosci Res, 95, 222–234.

Raymo, J.M. (2015) Living alone in Japan: Relationships with happiness and health. Demographic research, 32, 1267.

Ritchie, S.J., Cox, S.R., Shen, X., Lombardo, M.V., Reus, L.M., Alloza, C., Harris, M.A., Alderson, H.L., Hunter, S., Neilson, E., Liewald, D.C.M., Auyeung, B., Whalley, H.C., Lawrie, S.M., Gale, C.R., Bastin, M.E., McIntosh, A.M. & Deary, I.J. (2018) Sex Differences in the Adult Human Brain: Evidence from 5216 UK Biobank Participants. Cereb Cortex, 28, 2959–2975.

Robinson, G.E., Fernald, R.D. & Clayton, D.F. (2008) Genes and social behavior. Science, 322, 896–900.

Sallet, J., Mars, R.B., Noonan, M.P., Andersson, J.L., O’Reilly, J.X., Jbabdi, S., Croxson, P.L., Jenkinson, M., Miller, K.L. & Rushworth, M.F. (2011) Social network size affects neural circuits in macaques. Science, 334, 697–700.

Sallie, S.N., Ritou, V., Bowden-Jones, H. & Voon, V. (2020) Assessing international alcohol consumption patterns during isolation from the COVID-19 pandemic using an online survey: highlighting negative emotionality mechanisms. BMJ Open, 10, e044276.

Salvatier, J., Wiecki, T.V. & Fonnesback, C. (2016) Probabilistic programming in Python using PyMC3. PeerJ Computer Science, 2.

Sandford, A. (2020) Coronavirus: Half of humanity now on lockdown as 90 countries call for confinement.

Schaefer, A., Kong, R., Gordon, E.M., Laumann, T.O., Zuo, X.N., Holmes, A.J., Eickhoff, S.B. & Yeo, B.T.T. (2018) Local-Global Parcellation of the Human Cerebral Cortex from Intrinsic Functional Connectivity MRI. Cereb Cortex, 28, 3095–3114.

Schurz, M., Uddin, L., Kanske, P., Lamm, C., Sallet, J., Bernhardt, B., Mars, R. & Bzdok, D. (2021a) Variability in brain structure and function reflects lack of peer support. Cerebral Cortex.

Schurz, M., Uddin, L.Q., Kanske, P., Lamm, C., Sallet, J., Bernhardt, B.C., Mars, R.B. & Bzdok, D. (2021b) Variability in Brain Structure and Function Reflects Lack of Peer Support. Cereb Cortex.

Seo, H., Cai, X., Donahue, C.H. & Lee, D. (2014) Neural correlates of strategic reasoning during competitive games. Science, 346, 340–343.

Seo, H. & Lee, D. (2008) Cortical mechanisms for reinforcement learning in competitive games. Philos Trans R Soc Lond B Biol Sci, 363, 3845–3857.

Silk, J.B., Alberts, S.C. & Altmann, J. (2003) Social bonds of female baboons enhance infant survival. Science, 302, 1231–1234.

Smith, S.M., Zhang, Y., Jenkinson, M., Chen, J., Matthews, P.M., Federico, A. & De Stefano, N. (2002) Accurate, robust, and automated longitudinal and cross-sectional brain change analysis. Neuroimage, 17, 479–489.

Spreng, R.N., Dimas, E., Mwilambwe-Tshilobo, L., Dagher, A., Koellinger, P., Nave, G., Ong, A., Kernbach, J.M., Wiecki, T.V., Ge, T., Li, Y., Holmes, A.J., Yeo, B.T.T., Turner, G.R., Dunbar, R.I.M. & Bzdok, D. (2020) The default network of the human brain is associated with perceived social isolation. Nat Commun, 11, 6393.

Stahn, A.C., Gunga, H.C., Kohlberg, E., Gallinat, J., Dinges, D.F. & Kühn, S. (2019) Brain Changes in Response to Long Antarctic Expeditions. N Engl J Med, 381, 2273–2275.

Statistics, O.f.N. (2019) The cost of living alone.

Stefanacci, L. & Amaral, D.G. (2002) Some observations on cortical inputs to the macaque monkey amygdala: an anterograde tracing study. J Comp Neurol, 451, 301–323.

Stranahan, A.M., Khalil, D. & Gould, E. (2006) Social isolation delays the positive effects of running on adult neurogenesis. Nat Neurosci, 9, 526–533.

Tavares, R.M., Mendelsohn, A., Grossman, Y., Williams, C.H., Shapiro, M., Trope, Y. & Schiller, D. (2015) A Map for Social Navigation in the Human Brain. Neuron, 87, 231–243.

Von Der Heide, R., Vyas, G. & Olson, I.R. (2014) The social network-network: size is predicted by brain structure and function in the amygdala and paralimbic regions. Soc Cogn Affect Neurosci.

Vyas, A., Jadhav, S. & Chattarji, S. (2006) Prolonged behavioral stress enhances synaptic connectivity in the basolateral amygdala. Neuroscience, 143, 387–393.

Vyas, A., Mitra, R., Rao, B.S. & Chattarji, S. (2002) Chronic stress induces contrasting patterns of dendritic remodeling in hippocampal and amygdaloid neurons. Journal of Neuroscience, 22, 6810–6818.

Wang, H.-T., Smallwood, J., Mourao-Miranda, J., Xia, C.H., Satterthwaite, T.D., Bassett, D.S. & Bzdok, D. (2018) Finding the needle in high-dimensional haystack: A tutorial on canonical correlation analysis. arXiv preprint arXiv:1812.02598.

Watanabe, Y., Gould, E. & McEwen, B.S. (1992a) Stress induces atrophy of apical dendrites of hippocampal CA3 pyramidal neurons. Brain Res, 588, 341–345.

Watanabe, Y., Gould, E. & McEwen, B.S.J.B.r. (1992b) Stress induces atrophy of apical dendrites of hippocampal CA3 pyramidal neurons. 588, 341–345.

Weber, J., Javelle, F., Klein, T., Foitschik, T., Crucian, B., Schneider, S. & Abeln, V. (2019) Neurophysiological, neuropsychological, and cognitive effects of 30 days of isolation. Exp Brain Res, 237, 1563–1573.

Woolley, C.S., Gould, E. & McEwen, B.S. (1990a) Exposure to excess glucocorticoids alters dendritic morphology of adult hippocampal pyramidal neurons. Brain Res, 531, 225–231.

Woolley, C.S., Gould, E. & McEwen, B.S.J.B.r. (1990b) Exposure to excess glucocorticoids alters dendritic morphology of adult hippocampal pyramidal neurons. 531, 225–231.

Yuan, R., Nechvatal, J.M., Buckmaster, C.L., Ayash, S., Parker, K.J., Schatzberg, A.F., Lyons, D.M. & Menon, V. (2021) Long-term effects of intermittent early life stress on primate prefrontal-subcortical functional connectivity. Neuropsychopharmacology, 46, 1348–1356.

Yukie, M. (2002) Connections between the amygdala and auditory cortical areas in the macaque monkey. Neurosci Res, 42, 219–229.

Zhang, Y., Brady, M. & Smith, S. (2001) Segmentation of brain MR images through a hidden Markov random field model and the expectation-maximization algorithm. IEEE Trans Med Imaging, 20, 45–57.

